# Myeloid and endothelial cells cooperate to promote hematopoietic stem cells expansion in the fetal niche

**DOI:** 10.1101/2021.04.07.438809

**Authors:** Pietro Cacialli, Marie-Pierre Mailhe, Rachel Golub, Julien Y. Bertrand

## Abstract

During embryonic development, very few hematopoietic stem cells (HSCs) are produced from the hemogenic endothelium, that will be expanded in a very specific niche. This fetal HSC niche comprises a complex and dynamic molecular network of interactions between multiple cell types, including endothelial cells (ECs) and mesenchymal stromal cells. It is known that functional changes in the hematopoietic niche, such as aging, vascular cell remodelling or inflammation can directly affect HSCs. Among all these inflammatory regulators, the eicosanoid prostaglandin E (PGE2) has been shown to be very important during embryonic life. However, the precise source of PGE2 in the embryo is still elusive. Here we show that all the genes involved in PGE2 synthesis and transport are expressed by distinct cells of the caudal hematopoietic tissue (CHT) in the zebrafish embryo and in the mouse fetal liver, suggesting that each cell type acts sequentially and collaboratively with the others to produce PGE2 and ultimately expand HSCs. Among these cells, we found myeloid cells (both neutrophils and macrophages) to be absolutely necessary, as they concur to the production of PGH2, the precursor of PGE2. To measure the impact of myeloid cells, we generated a genetic model of myeloid ablation, which caused a loss of HSCs in the CHT, that could be rescued by supplementing zebrafish embryos with PGE2 or PGH2. ECs expressed the *slco2b1* transporter to import PGH2, and *ptges3*, the necessary enzyme to convert this latter into PGE2. Taken altogether, our data show that the triad composed of neutrophils, macrophages and ECs concurs to HSC expansion in the CHT.

## Introduction

Hematopoietic stem cells (HSCs) are a rare population endowed with self-renewal activity, and that can regenerate all blood lineages during fetal and adult life. The original pool of HSCs is established through many developmental processes that involve several specific microenvironments. These hematopoietic niches consist of different cell types, adhesion molecules and secreted/membrane-bound signalling factors that can directly affect HSCs or their progeny. The detailed understanding of the complex interactions between HSCs and their niche is therefore critical to improve HSC transplantation-based therapies. While *in vitro* culture systems have allowed a wide comprehension of key signalling involved in HSC differentiation, the mechanisms controlling HSC expansion, which only occurs during embryogenesis are still not fully appreciated (Lessard et al., 2004). The use of vertebrate model organisms such as mouse and zebrafish has contributed to clarify the mechanisms of hematopoietic development. HSCs first arise from the hemogenic endothelium of the dorsal aorta (DA) (Bertrand et al., 2010). Through blood circulation, they colonize a transient hematopoietic niche before they settle in the adult marrow. In mammals, this temporary niche is the fetal liver, whereas the caudal hematopoietic tissue (CHT) plays this role in zebrafish embryos (Tamplin et al., 2015). This transitory niche is the only one where HSCs will expand extensively, up to 40 times in the mouse embryo (Ema and Nakauchi, 2000). Due to the possibilities to manipulate the zebrafish embryo, we and others have discovered many signalling pathways involved in HSC expansion (Tamplin et al., 2015). Among these molecules, the prostaglandin E (PGE2) has shown promising results both experimentally and clinically (Goessling et al., 2011), although it was originally described as an enhancer of HSC specification from the hemogenic endothelium (North et al., 2007).

PGE2 is the most abundant prostaglandin in the organism. It is synthesized through the sequential oxygenation of arachidonic acid. Arachidonic acid derives from diacylglycerol or phospholipids, present in the cell membrane, that are oxidized by phospholipases (from the C or A2 family). Arachidonic acid is then metabolized by the cyclooxygenases *cox1* and *cox2* to produce PGG2/PGH2, before the *Prostaglandin-E-synthase (ptges)*, an isomerase, synthetises PGE2. Many studies showed that PGE2 enhances HSC specification and/or proliferation and in contrast, blocking prostaglandin synthesis, decreased HSC numbers (North et al., 2007). PGE2 can bind to four different G-coupled receptors, EP 1–4, which have various cellular functions (Sugimoto and Narumiya, 2007). Both EP2 and EP4 receptors are expressed by hematopoietic precursors (North et al., 2007), and transduce PGE2 signalling through the cAMP/PKA pathway. EP4 additionally activates the phosphatidylinositol 3-kinase (PI3K)/AKT pathway (Fujino et al., 2003). However, the precise cellular source of each PGE2 metabolite in the embryo has yet to be cleared. Here, we show that all the genes involved in PGE2 synthesis are expressed by different cells of the CHT in the embryonic zebrafish, a pattern that seems conserved also in the mouse fetal liver. In the zebrafish CHT, as in mouse fetal liver, we find that neutrophils express high levels of phospholipases, while macrophages express *cox1/2* enzymes and endothelial cells (ECs) high levels of *ptges*. This suggests that each cell type is sequentially necessary to mediate PGE2 synthesis. To measure the impact of myeloid cells, we generated a genetic model of myeloid ablation, which caused a loss of HSCs in the CHT, and could be rescued by supplementing zebrafish embryos with PGE2 or PGH2. Moreover, we identified the role of an important transporter, *slco2b1*, that mediates the transport of prostaglandins across the cell membrane into ECs. This transporter is a member of the Organic Anion Transporting Polypeptides (OATP) superfamily, that is largely conserved between zebrafish and mammals (Popovic et al., 2010). Indeed, all zebrafish OATPS consist of 12 transmembrane domains (TMD) with a larger fifth extracellular loop, LP5, that contains 10 conserved cysteine residues. These conserved cysteine residues within LP5 are found to be crucial for protein function (Hanggi et al., 2006). In the present report, we found a defect of HSC expansion in the CHT of *slco2b1*-deficient embryos, which could also be rescued by exogenous PGE2. Taken altogether, our data show that the myeloid cells and the vascular niche cooperate to enhance HSC expansion in the CHT.

## Results

### The PGE2 synthesis pathway is distributed over three different cell types in the CHT

In order to accurately describe the PGE2 synthesis pathway, we quantified the expression of the genes coding for the enzymes, channels and receptors involved in PGE2 signalling, in different populations of the zebrafish CHT. We dissected CHTs (caudal regions) at 48 hpf from *kdrl:GFP, ikaros:GFP, mpeg1:GFP, mpx:GFP* transgenic animals, and sorted GFP-positive cells to purify ECs, HSPCs, macrophages and neutrophils, respectively. We evaluated the expression of each of these genes by qPCR analysis (Fig.1a). We find that the phospholipases *pla2g4aa* and *pla2g4ab* are highly expressed in neutrophils (Fig.1b-c), while the expression of cyclooxygenases *Cox1, cox2a* and *cox2b* is enriched in macrophages (Fig.1d-e-f). Finally, we find that the prostaglandin synthases *ptges3a* and *ptges3b* are highly expressed in ECs (Fig.1g-h). As expected, and as previously reported (North et al., 2007) the prostaglandin receptors *ptger1a, ptger1b, ptger2a, ptger4a* are mostly expressed in HSPCs. *ptger3*, and *ptger4a* to a lesser extent, were expressed in neutrophils (Fig.1i-o). This data establishes that in the CHT, different cell types seem to play an important role in PGE2 production. PGE2 release and uptake are required to initiate and terminate their biological actions, respectively. Thus, efficient transport of PGE2 across the cell membrane is critical. This can be mediated by the transmembrane transporters ATP-binding cassette, subfamily C, member 4 (*abcc4*; also known as *MRP4*) and solute carrier organic anion transporter (OATP) family members, such as *SLCO2B1*. While *ABCC4* plays a role in PGE2 release from the producing cell (efflux PGE2 transporter) (Russel et al., 2008), *SLCO2B1* promotes PGE2 uptake from the extracellular space (influx PGE2 transporter) (International Transporter et al., 2010), but could also contribute to the uptake of other metabolites of the prostaglandin family (Li et al., 2020). Previous studies in mammals showed that *slco2b1* and *abcc4* are highly expressed in hepatocytes in the adult liver, but also in other tissues including the intestines, heart and brain capillary ECs (Tamai et al., 2000). In the zebrafish CHT, we find that *slco2b1* and *abcc4* prostaglandin transporters are highly expressed in caudal ECs during development, as determined by qPCR (Fig.1p-q), and confirmed by *in situ* hybridization at different stages of embryo development (Supp. Fig.1a-b). Altogether, our data suggest that PGE2 synthesis is achieved by many cell subsets in the hematopoietic niche.

**Figure 1.**
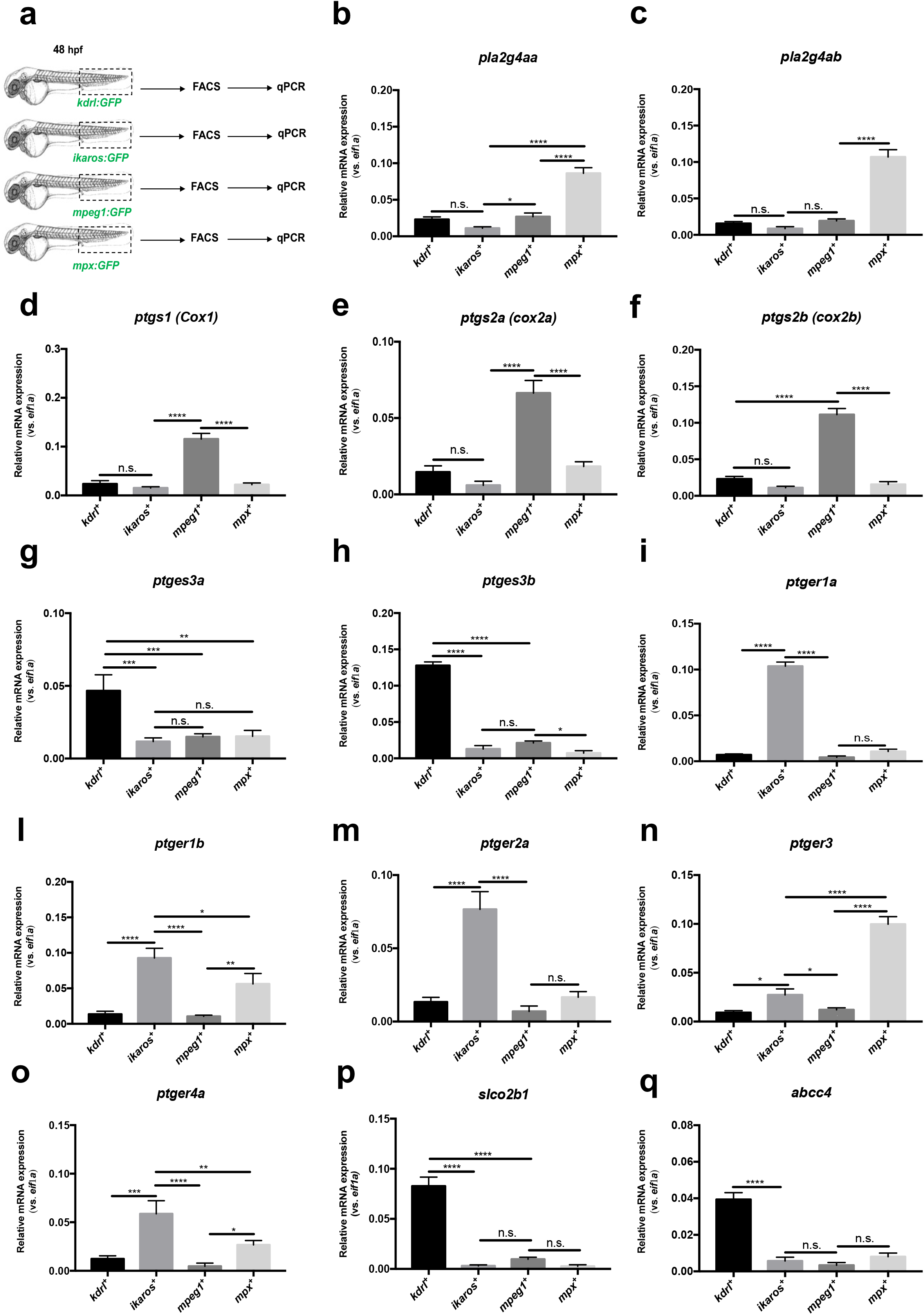
The PGE_2_ synthesis pathway in zebrafish CHT. (a) Experimental outline of qPCR analysis after dissection of CHT (caudal region) at 48 hpf from *kdrl:GFP, ikaros:GFP, mpeg1:GFP, mpx:GFP* transgenic animals, and FACS-sorted GFP-positive cells to purify ECs, HSPCs, macrophages and neutrophils. Data are biological triplicates plated in technical duplicates. (b-c) The phospholipases *pla2g4aa* and *pla2g4ab* are highly expressed in neutrophils. (d-e-f) The expression of cyclooxygenases *cox1, cox2a and cox2b* is enriched in macrophages. (g-h) The prostaglandin synthases *ptges3a* and *ptges3b* are only expressed by ECs. (i-o) The prostaglandin receptors *ptger1a, ptger1b, ptger2a, ptger3, ptger4a* are mostly expressed in HSPCs. (p-q) The prostaglandin transporters *slco2b1* and *abcc4* are specifically expressed in ECs. Statistical analysis was completed using one-way ANOVA, multiple comparison test. * P<.05; **P<.01; ***P<.001; ****P<.0001.

### The specific ablation of myeloid cell decreases the number of HSCs in the CHT

During zebrafish development, HSCs have been shown to mainly interact with ECs and stromal cells in the CHT (Tamplin et al., 2015). However, we show here that myeloid cells could have an important role as they express the key enzymes to initiate the production of PGE2. To investigate whether and how the myeloid cells contribute to HSPCs expansion, we genetically ablated these cell types during development. This was achieved using a *Tg(cd45:CFP-NTR)* zebrafish line in which the bacterial nitroreductase (NTR), fused to CFP, is expressed in *ptprc(cd45)*-positive cells (Suppl.Fig.2a). Although CD45 is a pan-leukocytic marker, we have previously shown in adult zebrafish that the 7.6kb promoter we isolated is only transcriptionally active in myeloid and T cells (Wittamer et al., 2011), but not in hematopoietic progenitors or any other hematopoietic subsets. To assess this during embryogenesis, we combined the *cd45:CFP-NTR* and *runx1:mCherry* transgenic lines and analysed double transgenic embryos at 72hpf by cytometry (Suppl.Fig.2b). Our data clearly show that CFP and mCherry do not overlap, meaning that the 7.6kb cd45 promoter is not active in embryonic HSPCs (Suppl.Fig.2c). As previously reported, the ablation system allows killing of NTR-expressing cells upon addition of metronidazole (MTZ) (Pisharath, 2007). Importantly, expression alone or administration of MTZ to non-transgenic embryos did not induce apoptosis. By contrast, a single treatment with MTZ between 48 and 72 hpf was sufficient to completely ablate myeloid cells in *cd45:CFP-NTR* transgenic embryos (Suppl.Fig.3a). Since our previous results suggested that myeloid cells have a relevant role in PGE2 synthesis in the CHT, we treated *cd45:CFP-NTR* transgenic embryos with DMSO and/or MTZ (between 48 and 72hpf), and found a significative decrease of PGE2 levels after myeloid ablation, as measured by ELISA (Suppl.Fig.3b), according to manufacturer’s protocols and as previously reported (Esain et al., 2015). This could be explained by the absence of phospholipases and cox enzymes after myeloid ablation, as scored by qPCR (Suppl.Fig.4a-f). Interestingly, after myeloid ablation, we found not change in the expression for the prostaglandin transporters *slco2b1, abcc4* and prostaglandin synthases *ptges3a* (Suppl.Fig.4g-h-i), but a small decrease of *ptges3b* (Suppl.Fig.4l). Finally, we find that myeloid ablation decreases the expression of prostaglandin receptors *ptger1a, ptger1b, ptger2a, ptger3, ptger4a* (Suppl.Fig.4m-n-o-p-q), which could be explained by the expression of these receptors by some myeloid subsets (Figure 1), but also probably by the impact on HSC expansion. These results confirm that myeloid cells have an important role in the production of PGE2 in the CHT niche, and that their absence cannot be compensated by other cells in the hematopoietic niche.

To verify this hypothesis, we combined the *cd45:CFP-NTR* and *runx1:mCherry* transgenic lines, and treated double transgenic embryos between 48 and 72hpf with DMSO and/or MTZ. As we expected, the number of *runx1:mCherry^+^* cells decreased in the CHT of double transgenic embryos at 72hpf, compared to controls (Fig.2a-b). We could rescue this phenotype by supplementing MTZ-treated double transgenic embryos with PGE2 and PGH2 (Figure 2a, b), but treatments with AA or PGG2 did not rescue the numbers of runx1-positive cells in the CHT (Suppl. Figure 5). We also quantified the number of *cd45*-positive cells and only PGE2 treatment could modestly rescue the number of myeloid cells after ablation (Fig.2c). In summary, our data indicate that myeloid cells are necessary for PGE2 synthesis, as they ultimately contribute to the production of PGH2, which will then need to be metabolised into PGE2 to promote HSC expansion.

**Figure 2.**
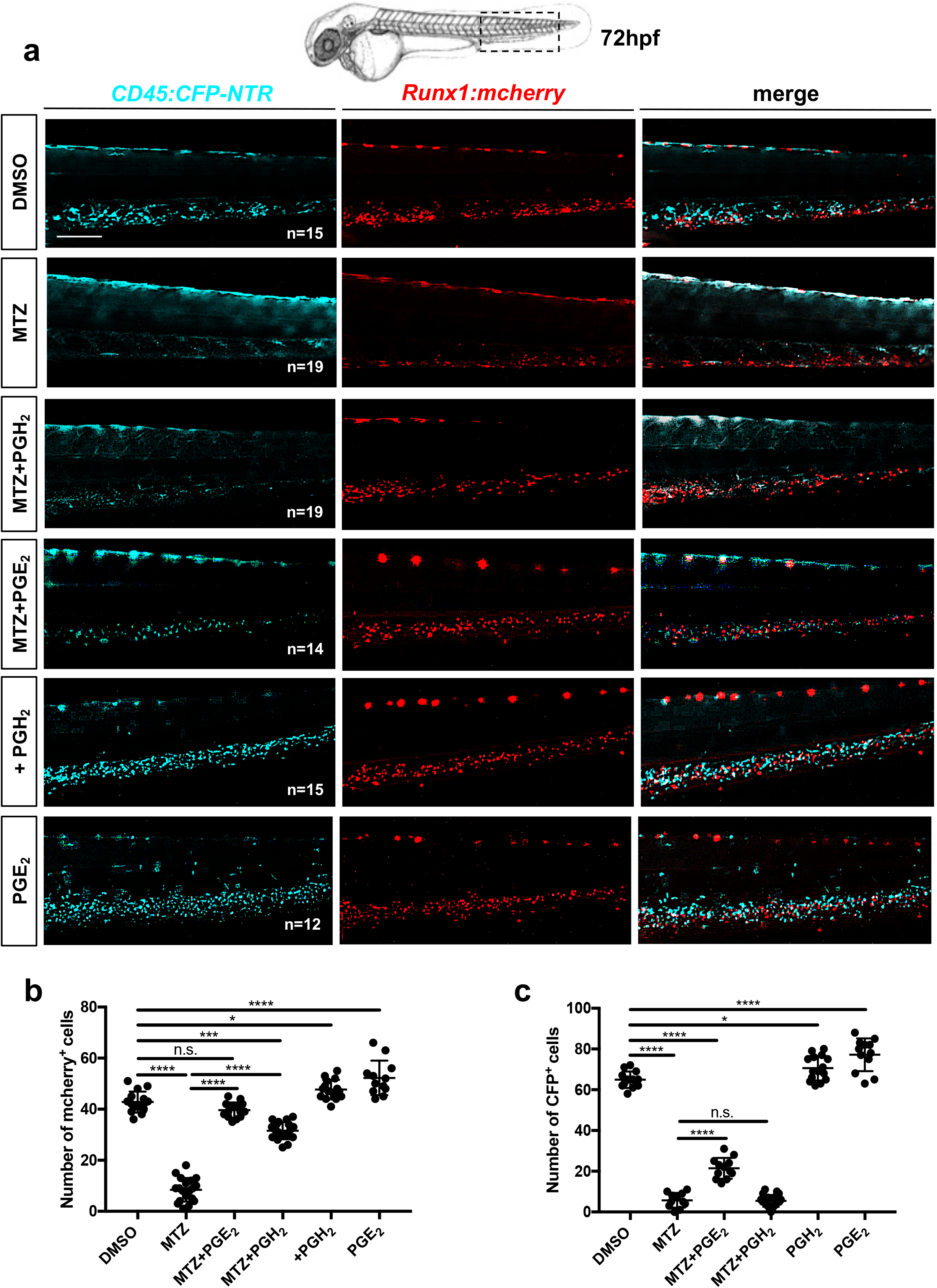
The loss of HSC after myeloid ablation can be rescued by PGE2 or PGH2 treatments. (a) Fluorescence imaging of the CHT of double transgenic *cd45:CFP-NTR/runx1:mcherry* embryos in DMSO and after treatment with MTZ and PGH2 or PGE2. (b) Quantification of *runx1:mcherry* positive-cells in double transgenic *cd45:CFP-NTR/runx1:mcherry* embryos in DMSO and after treatment with MTZ and PGH2 or PGE2. (c) Quantification of *cd45:CFP* positive-cells in double transgenic *cd45:CFP-NTR/runx1:mcherry* embryos in DMSO and after treatment with MTZ and PGH2 or PGE2. Statistical analysis was completed using one-way ANOVA, multiple comparison test. **P<.01; ****P<.0001. Scale bar 200μm (a).

### *slco2b1-deficient* embryos show a severe decrease in HSC expansion in the CHT

As mentioned before, myeloid cells can produce PGG2/PGH2 from arachidonic acid but are unable to convert this precursor into PGE2 as they lack the expression of *ptges* enzymes, that are however highly expressed in caudal ECs. Therefore, PGG2/PGH2 has to be transferred to ECs to be processed. OATP channels are specialized channels involved in intracellular influx (International Transporter et al., 2010). In particular, *SLCO2B1* has been shown to promote the entry of PGE2 or similar metabolites into different cell types. As *slco2b1* expression is highly enriched in ECs of the CHT, we then investigated the consequences of *slco2b1*-deficiency on HSCs during zebrafish embryo development. We used the uncharacterized *slco2b1^sa37367^* mutant line which presents a point mutation in the splice donor site at the end of exon 4 (Suppl.Fig.6a). By *in situ* hybridization, we show that *slco2b1*^-/-^ embryos exhibit a defect in definitive hematopoiesis as early as 60 hpf (Fig.3a-b) which was maintained at 4 dpf (Suppl.Fig.6b). However, no change in *runx1* and *cmyb* expression were detected at 28hpf and 36hpf, respectively, showing that *slco2b1* is not involved in HSC specification from the hemogenic endothelium (Suppl.Fig.6c-d). Of note, *slco2b1*-deficient embryos did not show any changes in the development of early vasculature or in primitive hematopoiesis (Suppl.Fig.7a-b-c), showing an exclusive role for *slco2b1* during HSPC expansion in the CHT. Morpholino-mediated knockdown of *slco2b1* expression completely phenocopied *slco2b1*^-/-^ mutant embryos, where HSC specification was unaffected but their expansion impaired in the CHT (Suppl.Fig.8a-b-c). To confirm our results, we first injected control- and *slco2b1*-morpholinos in *kdrl:mcherry;cmyb:GFP* embryos and scored the number of HSPCs at 60 hpf, where we observed a significant decrease of double-positive cells in *slco2b1*-morphants (Fig.3c-d). Next, we used time-lapse confocal imaging to follow the HSCs’ behaviour in the CHT in *cmyb:GFP slco2b1*-morphants. While the number of HSCs augmented in control embryos between 54 and 60 hpf, their number remained unchanged in *slco2b1*-deficient embryos (Fig.3e and Suppl. videos 1-2). These results suggested that the absence of *slco2b1* in the vascular niche generated a defect of HSPC proliferation in the CHT. To confirm this hypothesis, we injected control- and *slco2b1*-morpholinos in *cmyb:GFP* embryos and stained for both GFP and phospho-Histone 3 (pH3) to quantify proliferating HSPCs. At 60 hpf, *slco2b1*-morphants showed a significant decrease in the number of proliferating HSPCs (Fig.4 a-b). Therefore, the function of *slco2b1* in ECs is necessary to the expansion of HSCs in the CHT.

**Figure 3.**
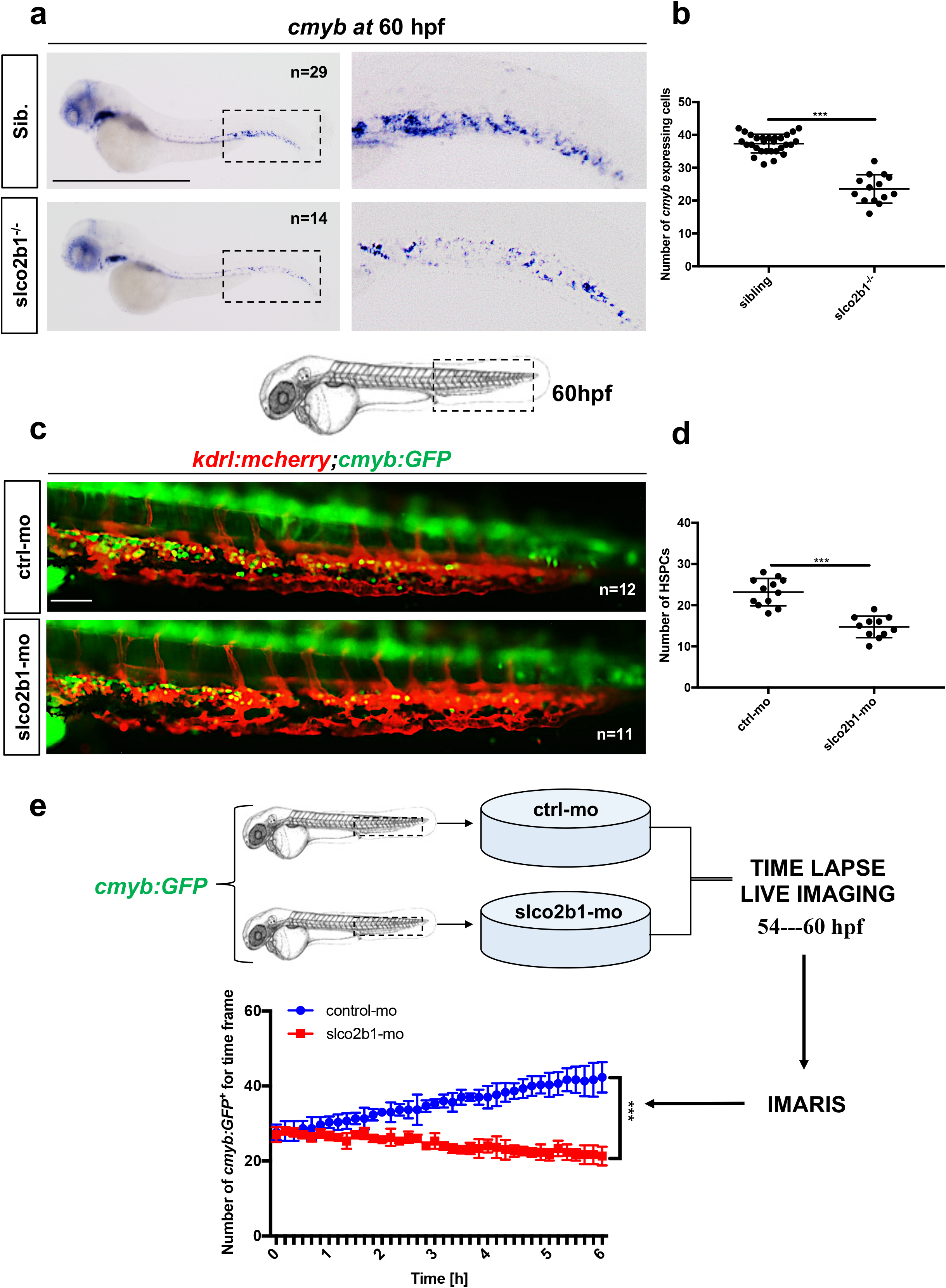
The deficiency of *slco2b1* induces a decrease of HSPC in the CHT. (a) WISH for *cmyb* expression at 60hpf in wild type and *slco2b1*^-/-^ embryos. (b) Quantification of cmyb-expressing cells. Statistical analysis: un-paired two tailed t test, ***P <.001. (c) Fluorescence imaging in the CHT of *kdrl:mCherry;cmyb:GFP* embryos injected with control- and *slco2b1-MOs* (d) Quantification of HSPCs associated to ECs. Statistical analysis: un-paired two tailed t test, ***P <.001. (e) Experimental outline and quantification of time-lapse live imaging in controls and slco2b1-morphants *cmyb:GFP*^+^ cells, using Imaris software. Centre values denote the mean, and error values denote s.e.m. The statistical analysis was completed using an unpaired two tailed t test. ***P < .001 Scale bar 500μm (a); 200μm (c)

**Figure 4.**
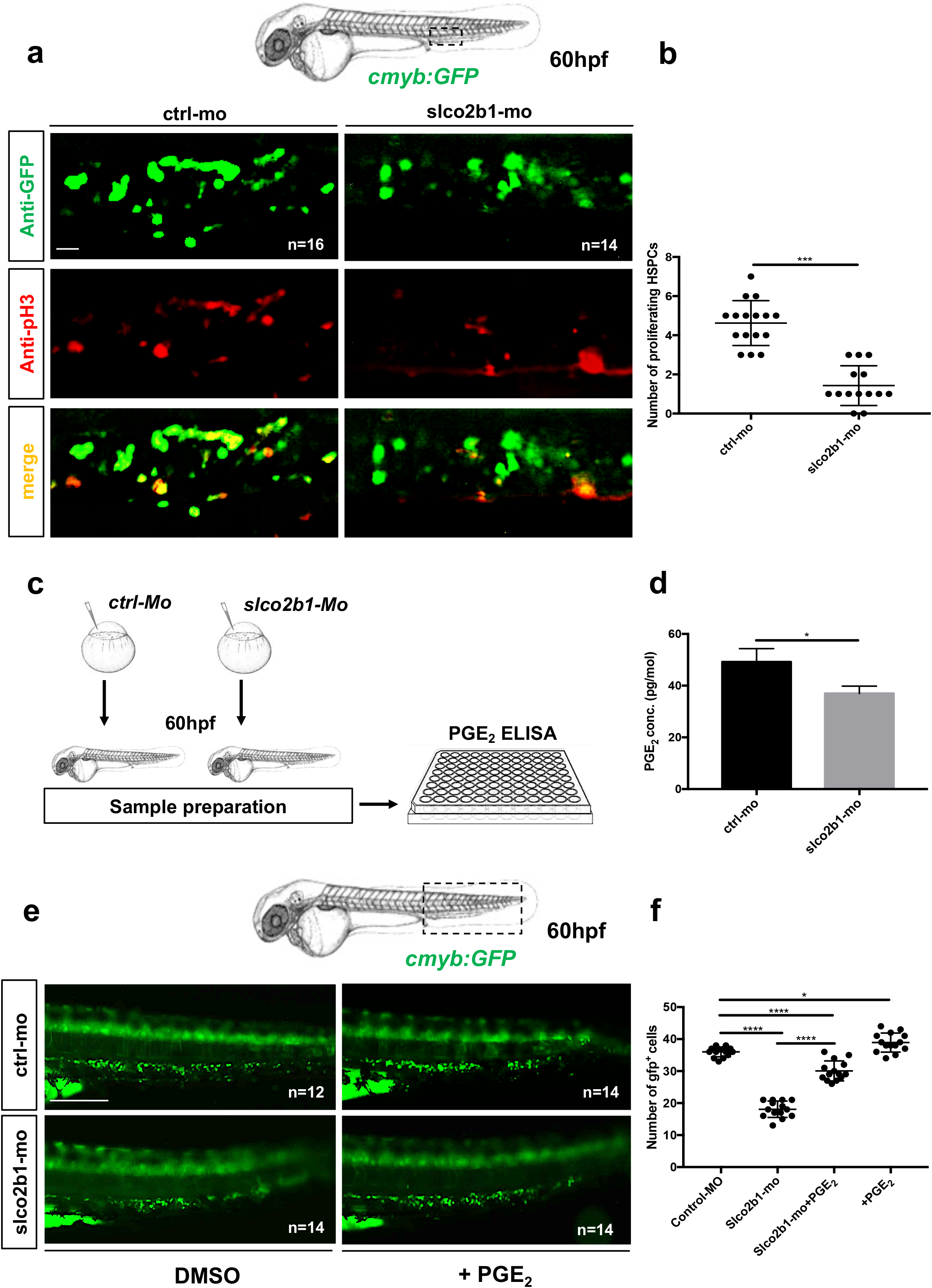
The defect of HSPCs proliferation in *slco2b1*-deficient embryos can be rescued by PGE2 treatment. (a) Anti-GFP and pH3 immunostainings of either controls (Ctrl-Mo) or *slco2b1*-morphants *(slco2b1-MO) cmyb:GFP* embryos. (b) Quantification of the number of the pH3+ HSCs in controls or *slco2b1*-morphants. Centre values denote the mean, and error values denote s.e.m, statistical analysis was completed using an unpaired two tailed t test. ***P < .001. (c) Experimental outline to measure PGE2 level by ELISA-kit in control and *slco2b1-MO* at 60hpf. (d) Quantification of PGE2 concentration in control and *slco2b1* morphants. The statistical analysis was completed using an unpaired two tailed t test *P < .01. (e) Fluorescence imaging in the CHT of *cmyb:GFP* embryos injected with control- and *slco2b1* - MOs and treated with PGE2. (f) Quantification of *GFP* positive cells. Statistical analysis: oneway ANOVA, multiple comparison, *P < .01; ****P < .0001. Scale bar 50μm (a); 200μm (e)

### PGE2, but not PGH2, rescues the loss of HSCs observed in *slco2b1*-deficient embryos

Based on our results, the deficiency of *slco2b1* induces a defect of HSPC proliferation, probably caused by a decrease of PGE2 in the CHT of zebrafish embryo. However, as mentioned earlier, *slco2b1* is a transporter used for the uptake of PGE2, and other prostaglandins, therefore the absence of *slco2b1* could potentially result in an increase of PGE2 in the extracellular environment. In order to evaluate the amount of PGE2 in *slco2b1*-morphants, we injected wild type AB* embryos with control- and *slco2b1*-morpholinos, and at 60hpf we processed embryos and measured PGE2 levels by ELISA assay (Fig.4c-d). We found a significant decrease of PGE2 level in *slco2b1*-morphants compared with controls (Fig.4d), suggesting that less PGE2 is produced in these embryos, concordant with the defect observed in HSPCs. Therefore, we investigated the role of *slco2b1* in PGE2 synthesis. First, we injected AB* embryos with control- and *slco2b1*-morpholinos and supplemented them with PGE2, PGH2, PGG2 or AA, from 48 to 60hpf. We then anlaysed *cmyb* expression by WISH and counted the number of *cmyb*-expressing cells in the CHT at 60hpf (Suppl.Fig.9a). We found that only PGE2 treatment rescued the loss of HSPCs in *slco2b1*-morphants (Suppl.Fig.9b). This was confirmed by live imaging using the *cmyb:GFP* transgenic reporter (Fig. 4e-f), and we also confirmed that HSPCs numbered where similarly rescued in *slco2b1* mutant embryos (Suppl.Fig.9c-d). As PGH2 (normally produced by macrophages) could not rescue the *slco2b1* deficiency, we hypothesised that ECs could import PGH2 through this transporter. To verify this, we injected *cd45:CFP-NTR;runx1:mCherry* embryos with control- and *slco2b1*-morpholinos and we treated with DMSO, MTZ and/or PGH2, from 48 to 72hpf, and counted the number of *mCherry^+^* cells at 72hpf for all conditions. As expected, *slco2b1*-morphants treated with DMSO showed less HSPCs (mCherry^+^) than control embryos (Fig.5a-d). Control- and *slco2b1*-morphants embryos treated with MTZ, where myeloid cells (CFP^+^) were ablated, showed a strong decrease of HSPCs (Fig.5b-d). Interestingly, control-morphants treated with MTZ and supplemented with PGH2 showed fairly normal numbers of HSPCs, in spite of the ablation of most of myeloid cells, but MTZ-treated *slco2b1*-morphants were not rescued by PGH2 (Fig.5c-d). We also quantified the number of *cd45*-positive cells and PGH2 could not rescue the number of myeloid cells after ablation (Fig.5e). This data confirms the hypothesis that in the absence of *slco2b1* transporter, caudal ECs cannot import the myeloid-derived PGH2, a metabolite necessary to the production of PGE2. Taken all together, our data suggests that the function of *slco2b1* in ECs is necessary for the expansion of HSPCs. In order to investigate the specific role of *slco2b1* in ECs and/or macrophages, we overexpressed *slco2b1* transiently by injecting the Tol2-UAS:*slco2b1* construct together with *tol2* mRNA in either *kdrl:Gal4* or *mpeg1:Gal4* embryos, and we performed WISH for *slco2b1* to confirm integration and tissue-specific overexpression (Suppl.Fig.10a-b). Next, to evaluate the impact of *slco2b1*-overexpression in ECs and/or macrophages on HSPCs, we scored *cmyb* at 60hpf in Gal4-reporter embryos injected with control- or *slco2b1*-morpholinos and/or co-injected with Tol2-UAS:*slco2b1* and *tol2* mRNA. As the slco2b1-morpholino targets a splice junction (Suppl.Fig.8), this experiment allowed us to selectively rescue *slco2b1* expression in ECs or macrophages, when the endogenous expression of *slco2b1* was abolished in all cell types. We found that the deficiency of HSPCs in *slco2b1*-morphants can be rescued only when slco2b1 expression was restored into caudal ECs (Suppl.Fig.11a-b-c), but not in macrophages (Suppl.Fig.12a-b-c). These results confirm that *slco2b1* transporter has a specific role in CHT-ECs by promoting the import of macrophage-derived PGH2 to synthetize PGE2.

**Figure 5.**
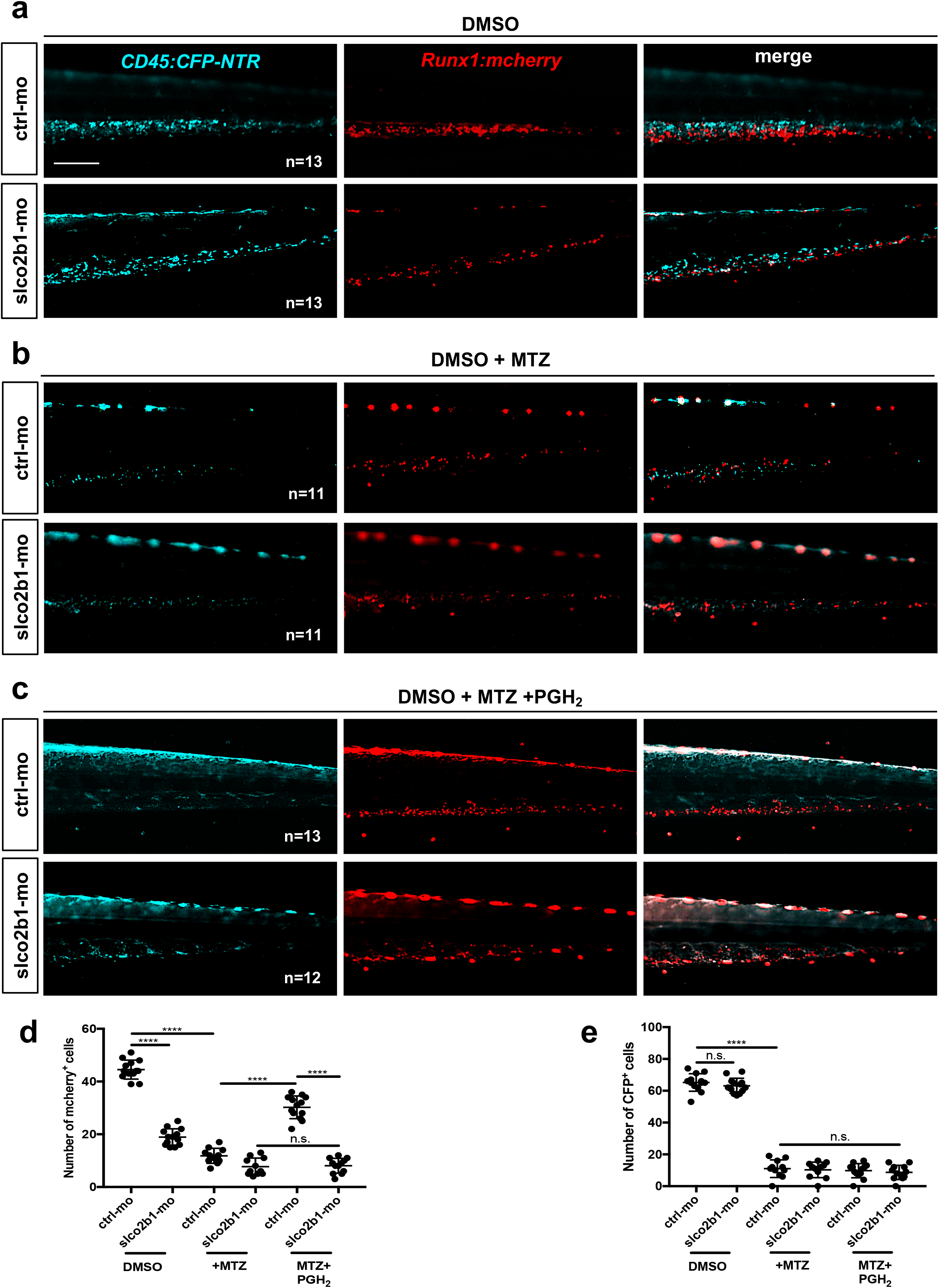
*Slco2b1* is necessary to import PGH2 into ECs. (a) Fluorescence imaging of the CHT of double transgenic *cd45:CFP-NTR/runx1:mcherry* embryos injected with control- or *slco2b1*-morpholinos in DMSO, after treatment with MTZ and PGH2. (b) Quantification of *runx1:mcherry* positive-cells. (c) Quantification of *cd45:CFP* positive-cells. Statistical analysis was completed using one-way ANOVA, multiple comparison test. ****P<.0001. Scale bar 200μm (a).

### The PGE2 synthesis pathway is conserved in the mouse fetal liver

In order to determine if the prostaglandin pathway present in the zebrafish CHT was also conserved in mammals, we performed qPCR on FACS-sorted cell subsets from mouse fetal liver at Embryonic day (E) 14.5 by using various combinations of antibodies. HSCs were isolated based on their Lin^-^ Sca1^+^cKit^+^ (LSK) phenotype; F4/80 also known as EMR1 or Ly71, was used as a macrophage marker, *Flk1* and CD31 to mark ECs and Gr-1, known as Ly-6G/Ly-6C, to sort neutrophils. By qPCR, we find that the phospholipase *pla2g4a* was highly expressed in neutrophils (Suppl.Fig.13a), while *pla2g4b* and *pla2g4c* were expressed at very low levels (Suppl. Figure 13b-c). As in the zebrafish, the cyclooxygenases *Cox1* and *Cox2* were mostly expressed in macrophages (Suppl.Fig.13d-e). We also find a high enrichment of prostaglandin synthase *Ptges3, slco2b1* and *Abcc4* in ECs (Suppl.Fig.13f-h). Next, we examined the expression of prostaglandin receptors *Ptger1, Ptger2, Ptger3, Ptger4* which were mostly expressed in HSCs, except for the receptor *Ptger2* that was only expressed in macrophages (Suppl.Fig.13i-n). By comparing zebrafish and mouse patterns of expression, we confirm the conservation of the prostaglandin pathway for each cell type between the zebrafish CHT and the mouse fetal liver (Fig.6a-b). Based on these results, we propose a model in which several cell types in the CHT niche, and in the mouse fetal liver, concur together to the production of PGE2, and therefore to the expansion of HSCs (Fig.6c-d).

**Figure 6.**
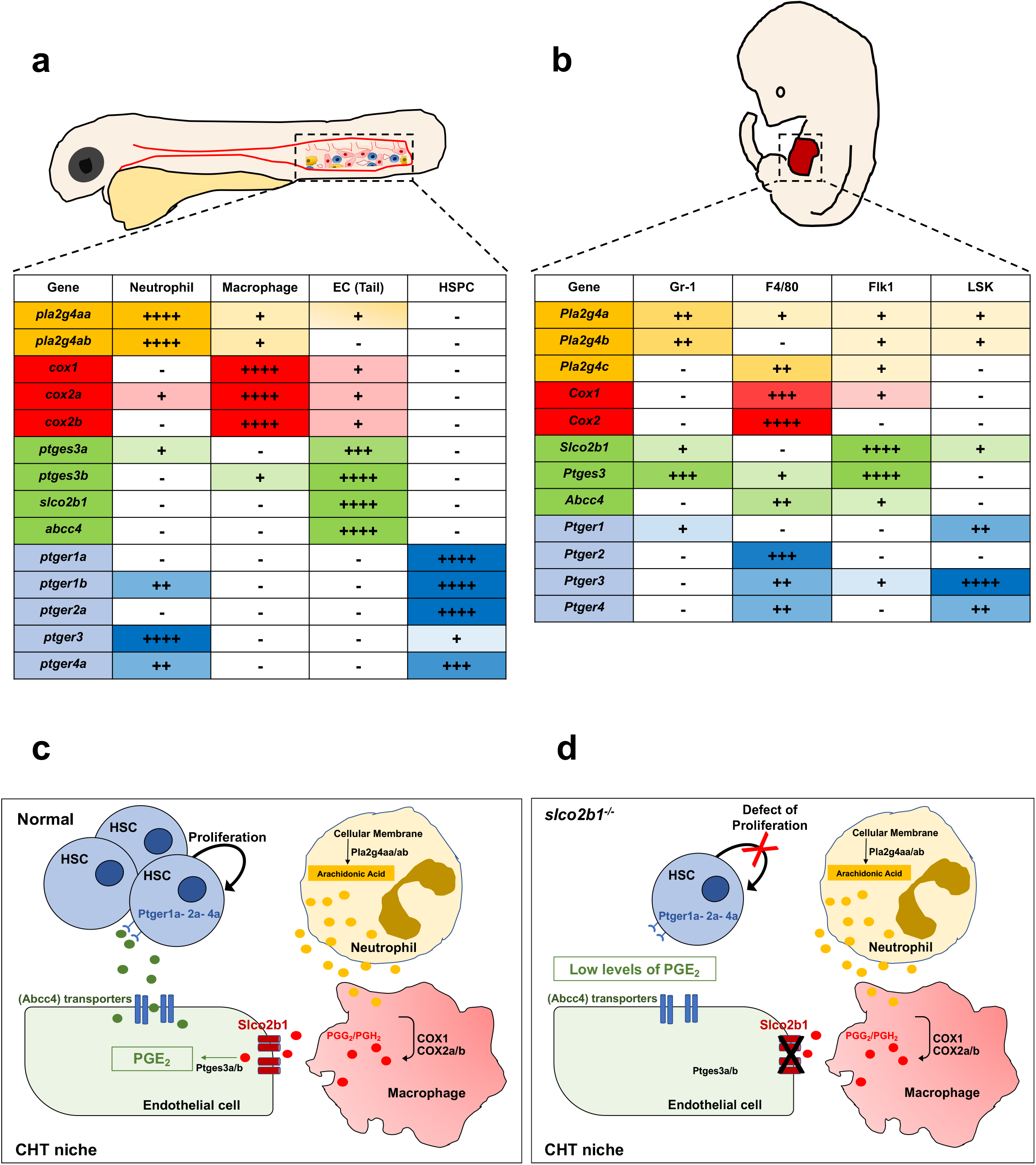
The prostaglandin synthesis pathway is conserved between the zebrafish CHT and the mouse fetal liver. (a) Summary of prostaglandin gene expression pathways in the CHT of zebrafish, and (b) in mouse fetal liver. (c) Proposal mechanism in which, HSCs, ECs, macrophages and neutrophils cooperate in the embryonic niche. In normal conditions *slco2b1* permits the transfer of PGE2 and precursor in ECs. (d) The deficiency of *slco2b1* decreases PGE2 levels in the CHT niche, generating a defect of HSC proliferation.

## Discussion

PGE2 plays important roles during inflammation and the immune response. However, these effects can be opposite depending on the cell type targeted. Indeed, different immune cell types will express different PGE2 receptors. These receptors belong to the GPCR family and will therefore transduce the PGE2 signal via a Gαq protein (EP1), a Gαs protein (excitatory receptors EP2 and EP4) or through a Gαi protein (inhibitory receptor EP3) (North et al., 2007). Indeed, many studies have shown that PGE2 not only stimulates the specification of HSCs from their immediate precursor, the hemogenic endothelium (North et al., 2007), but also their expansion, which has been successfully translated to non-human primates and patients (Goessling et al., 2011). The mammalian fetal liver, or the CHT in the zebrafish embryo, are thought to be the only niches where HSPCs actively proliferate to expand the population that was generated in the aorta. Therefore, we hypothesized that the PGE2 synthesis pathway was active in this microenvironment. Surprisingly, we found that all the key enzymes necessary to the synthesis of PGE2 (phospholipases, cox1/2 and prostaglandin-E-synthase) are distributed between different cell types: the neutrophils, macrophages and ECs. This finding involves a close collaboration between these cell types, and also involves that metabolites can transit from one cell type to another. We actually showed here that *Slco2b1* was important as an influx channel to import PGH2 into ECs, so that they can produce PGE2. Whereas, all neutrophils and macrophages might express the necessary enzymes, we found that *slco2b1* expression was restricted to ECs in the CHT. Moreover, when *slco2b1*-morphants showed a strong decrease of HSPCs numbers, restoring *slco2b1* in ECs was the only way to rescue HSPCs numbers, whereas overexpression in macrophages did not rescue their numbers. *abcc4*, the efflux transporter that releases PGE2 in the microenvironment, was also specifically expressed by ECs. Therefore, we can conclude that between 48 and 60 hpf, CHT-ECs are the sole source of PGE2 for expanding HSPCs, a role that is not surprising as CHT-ECs also produce other cytokines controlling HSPC expansion such as *kitlgb* (Mahony et al., 2016) and *osm* (Mahony et al., 2018).

During embryogenesis, many studies have shown the importance of primitive myeloid cells during the endothelium-to-hematopoietic transition. Indeed, macrophages seem to play many different roles: aortic macrophages are responsible, together with neutrophils, to create a pro-inflammatory environment that promotes HSC specification from the hemogenic endothelium, as well as a first round of expansion in the aortic niche (Espin-Palazon et al., 2014). Macrophages are then responsible for the emigration of HSCs out of the subaortic environment, as they can remove the extra-cellular matrix to allow newly born HSCs to enter intravasate in the cardinal vein (Travnickova et al., 2015), an effect that could be mediated through metalloproteases (Theodore et al., 2017). Finally, a recent study provided new insights into the mechanism of HSPC homing and reveals the essential role of VCAM-1-positive macrophages in HSPC retention in the CHT (Li et al., 2018). These macrophages, as well as the ones responsible for PGH2 production also likely come from the primitive, but not from the definitive wave, as recently demonstrated by fatemapping (Ferrero et al., 2021).

It has also been known for a long time, that macrophages, in mammals, contribute actively to adult hematopoiesis, either by helping red blood cells to differentiate in erythroblastic islands (Rhodes et al., 2008) or to remove old erythrocytes in the red pulp of the spleen (Nagelkerke et al., 2018). Similar processes have been observed in the fetal liver and fetal spleen in the mammalian embryo (Bertrand et al., 2006). In the bone marrow, α-SMA-positive macrophages interact tightly with HSPCs. They express high levels of *Cox2* which function seems to be important to maintain the long-term repopulation capability of HSCs, through the production of PGE2 (Ludin et al., 2012), a mechanism that seems similar to what we observe. However, in this study, the expression of *Ptges*, the enzyme that synthetizes PGE2 was not examined, and a contribution from the vascular niche cannot be excluded. Finally, it was recently shown that neutrophils are important for the hematopoietic recovery after irradiation. By irradiating neutropenic animals, Ballesteros and colleagues could show that the recovery of most hematopoietic lineages was delayed (Ballesteros et al., 2020). All these evidences, together with our observation that mouse fetal liver cells express the PGE2 pathway enzymes similarly to the zebrafish CHT populations, point to the existence of a triad, composed of neutrophils, macrophages and ECs, that controls the fate of HSPCs in all vertebrates.

## Acknowledgements

We would like to thank all lab members for comments. We are grateful to A. Cumano and S. Meunier for their help in sorting mouse fetal liver subsets. RG work is supported by Institut Pasteur, Institut National de la Santé et de la Recherche Médicale, Université de Paris, and ANR project NASHILCCD8 (#18-CE15-0024-01). JYB is funded by the Swiss National Fund (310030_184814).

## Author contributions

P.C. performed all zebrafish experiments. M.P.M. and R.G. performed experiments in the mouse embryo. R.G. edited the manuscript. P.C. and J.Y.B. designed experiments, performed analysis and wrote the manuscript.

## Raw data repository

All raw data is freely accessible on the following link: TBD / Yareta repository database.

## Material and method

### Zebrafish husbandry

AB*zebrafish strains, along with transgenic strains and mutant strains, were kept in a 14/10 h light/dark cycle at 28°C. Embryos were obtained as described previously. In this study we used the following transgenic lines: *Tg(mpeg1:GFP)^gl22^* (Ellett et al., 2011); *Tg(Mmu.Runx1:NLS-mCherry)^cz2010^* (here denoted as *runx1:mCherry*) (Tamplin et al., 2015); *Tg(cmyb:GFP)^zf169^* (North et al., 2007)*; Tg(kdrl:Has.HRASmCherry)^s896^* (Chi et al., 2008); *Tg(kdrl:GFP)^s843^* (Jin et al., 2005); *Tg(mpx:EGFP)^i113^* (Ellett et al., 2011); *Tg(kdrl:Gal4)^bw9^* (Kim et al., 2014); *Tg(mpeg1:Gal4)^g124^* (Ellett et al., 2011) *and Tg*(*cd45:CFP-NTR*).

### Cell sorting and flow cytometry on zebrafish embryos

Transgenic zebrafish embryos were incubated with a Liberase-Blendzyme 3 (Roche) solution for 90 min at 33°C, then dissociated and resuspended in 0.9× PBS-1% fetal calf serum, as previously described. We excluded dead cells by SYTOX-red (Life Technologies) staining. Cell sorting was performed using an Aria II (BD Biosciences).

### Quantitative real-time PCR and analysis on sorted cell by zebrafish

Total RNA was extracted using RNeasy minikit (Qiagen) and reverse transcribed into cDNA using a Superscript III kit (Invitrogen). Quantitative real-time PCR (qPCR) was performed using KAPA SYBR FAST Universal qPCR Kit (KAPA BIOSYSTEMS) and run on a CFX connect real-time system (Bio Rad). All Primers used are listed in Supplemental Table1.

### Generation of transgenic animals

The *Tg*(*cd45:CFP-NTR*) line was created using the Tol2 transposon system (Urasaki et al., 2006). Targeted cell ablation mediated by bacterial nitroreductase (NTR) was described previously (Curado et al., 2007). A DNA fragment containing CFP-NTR was subcloned into a Tol2 vector that contained the zebrafish cd45 promoter. Although CD45 is a pan-leukocytic marker, we have previously shown that the 7.6kb promoter we isolated is only reporting activity in myeloid and T cells (Wittamer et al., 2011), but not in HSPCs. The Tol2 construct and transposase mRNA were microinjected into 1- to 4-cell stage embryos and founders were screened based on the presence of CFP in myeloid and T cells in the progeny.

### Chemical treatments and analysis

The concentrations of all chemical treatments were chosen based on previous studies which established that these doses produced sub-lethal effects in zebrafish embryos, with no gross malformations (North et al., 2007). All compounds used in these experiments were purchased from Sigma-Aldrich. Zebrafish embryos were exposed for 12 hours as control in 0.2% dimethyl sulfoxide (DMSO), or 10μm of PGE_2_ starting at ~48hpf. For metronidazole (MTZ) treatment (for myeloid ablation) the *Tg(cd45:CFP-NTR)* embryos were exposed to 10mM of MTZ in 0.2% DMSO, from 48hpf to 72hpf in embryo. After exposure, fish were fixed and the expression of different markers was tested by WISH. Embryos were phenotyped based on the expression of the markers and eventually genotyped (when treating *slco2b1* mutants).

### Identification of *slco2b1* mutant line

The *slco2b1^sa37367^* mutant line which presents a point mutation T-C in splice donor site at the end of exon 4. The *slco2b1^sa37367^* line is available at the ZFIN repository (ZFIN ID: ZDB-ALT-160601-5326) from the Zebrafish International Resource Center (ZIRC). The *slco2b1^sa37367^* used in this paper was subsequently outcrossed with WT AB for clearing of potential background derived from the random ENU mutagenesis from which this line was originated. The *slco2b1^sa37367^* homozygous mutant is referred in the paper as *slco2b1*^-/-^ and was obtained by incrossing our *slco2b1^sa37367^* / AB strain. Genotyping was performed by PCR of the *slco2b1* gene followed by sequencing. Genotyping primers are: slco2b1-F: ATACCAGACTCAACTCCAGC; slco2b1-R:

TGTCTATGTCGACATACAAGC.

### Morpholino injections

The *slco2b1*-morpholino oligonucleotide (MOs) and control-MO were purchased from GeneTools (Philomath, OR). MO efficiency was tested by reverse transcription polymerase chain reaction (RT-PCR) from total RNA extracted from ~10 embryos at 48hpf. In all experiments, 12ng of *slco2b1*-MO were injected per embryo. Morpholinos and primer sequences are:

*Standard control-MO* CCTCTTACCTCAGTTACAATTTATA;
*slco2b1-MO* CATCCATGCTTTTTATCCTTGCCTC;
*slco2b1-Forward* CTCCAGCTCTTCAGTCTCAG;
*slco2b1-Reverse* CTGTGTCTGGCAAAGGCT.

### Transient overexpression of *slco2b1* in ECs and/or macrophage

For the *Tol2-UAS:slco2b1* construct, a Tol2 vector containing 4xUAS promoter, the sequence for *slco2b1* (including STOP codon), and a poly-adenylation signal sequence was generated by sub-cloning. Next, to overexpress *slco2b1* in ECs and/or macrophage, *Tg(kdrl:Gal4)* or *Tg(mpeg1:Gal4]* embryos were injected with 25pg of the final *Tol2-UAS:slco2b1* vector, and with 25pg *tol2* transposase mRNA. We performed WISH for *slco2b1* to confirm the integration of the Tol2 construct, and WISH for *cmyb* to evaluate the impact on HSPCs. We used the following primers to clone the full-length *slco2b1* coding sequence: *Full-Slco2b1-Forward* AAA GGA TCC CTC CAG CTC TTC AGT CTC AG; *Full-Slco2b1-Reverse* AAA CTC GAG TGG CCT GTA CAA CTG CTT GC.

### Whole-mount in situ hybridization and analysis

Digoxigenin and fluorescein-labeled probes *cmyb, runx1, pu1, gata1, flk1*, and *rag1* were previously described. Whole-mount in situ hybridization (WISH) was performed on 4% paraformaldehyde-fixed embryos. All injections were repeated 3 separate times. Analysis was performed using the un-paired Student t-test or ANOVA Multiple comparison-test (GraphPad Prism). Embryos were imaged in 100% glycerol, using an Olympus MVX10 microscope. Oligonucleotide primers used to amplify and clone cDNA for the production of the *slco2b1* and *abcc4* ISH probe are: *slco2b1*-F TGGTTGGGATTCCTGATAGC; *slco2b1*-R GGAAAGTGAAGCCACAAAGC; *abcc4*-F CCAGTCGACCTTCAGGA: *abcc4*-R CAGGAACAGGAAGCAAATCAAC.

### Confocal microscopy and immunofluorescence staining

Transgenic fluorescent embryos were embedded in 1% agarose in a glass-bottom dish. Immunofluorescence double staining was performed as described previously, with chicken anti-GFP (1:400; Life Technologies) and rabbit anti-phospho-histone 3 (pH3) antibodies (1:250; Abcam). AlexaFluor488-conjugated anti-chicken secondary antibody (1:1000; Life Technologies) and AlexaFluor594-conjugated anti-rabbit secondary antibody (1:1000; Life Technologies) to reveal primary antibodies. Confocal imaging was performed using a Nikon inverted A1r spectral.

### Time-lapse imaging and analysis

For time-lapse imaging, *Tg(cmyb:GFP*) embryos were anaesthetized with 0.03% Tricaine (Sigma), and embedded in 1% agarose in a 60mm glass-bottom dish. The embryos were imaged at 28.5□°C with a confocal Nikon inverted A1r spectral. Scanning with 20x water immersion objective, z-stalks were acquired with a step size of 7μm within an interval of 10 min for 6 hours in the control and *slco2b1*-morphants starting at ~54hpf. The analysis of *cmyb:GFP*^+^ cells in all experiments was performed using IMARIS image software.

### Enzyme-Linked Immunosorbent Assay (ELISA) for PGE2

ELISA for PGE2 was conducted using the Prostaglandin E2 Human ELISA Kit (Invitrogen - Thermo Fisher Scientific) according to manufacturer’s instructions. 50 embryos *cd45-NTR:CFP* treated with DMSO or MTZ were collected at 72hpf. 50 embryos AB* injected by control and *slco2b1*-morpholino were collected at 60hpf, washed in ice-cold PBS, and homogenized in 500μl TRIS buffer. Homogenate was spun down at 1200□rpm for four min at 4□°C to eliminate particulate, and supernatant collected for ELISA. Assays were run in technical triplicate.

### Mice

C57BL/6 mice were purchased from Charles River. Mice were cared for in accordance with Pasteur Institute guidelines in compliance with European animal welfare regulations; all animal studies were approved by Pasteur Institute Safety Committee according to the ethic charter approved by French Agriculture ministry and to the European Parliament Directive 2010/63/EU.

### Fetal liver cell preparations

E14.5 fetal livers were harvested, dissociated and resuspended in Hanks’ balanced-salt solution (HBSS) supplemented with 1% FCS (Gibco). Fetal liver cells were depleted of Lineage^+^ cells by staining with biotinylated-conjugated antibodies to lineage markers CD19 and Ter119 followed by incubation with streptavidin microbeads (Miltenyi Biotec). Depletion were performed on LS+MACS columns (Miltenyi Biotec), from which the negative fraction was recovered and stained for cell sorting.

### Fetal liver cell sorting

Flow cytometry data were analysed with FlowJo software (TreeStar). Dead cells were eliminated by Dead-Live staining (Thermo Fischer) exclusion. Fetal liver subsets were purified by sorting with a FACS Aria III (Becton Dickinson). Cells were recovered in eppendorf tubes for gene expression analyses.

All antibodies were from BD Biosciences, eBioscience, Biolegend, Sony, Cell signaling Technologies or R&D Systems. Antibodies either biotinylated or conjugated to fluorochromes (FITC, PE, PECy7, APC, APCCy7, BV510) were used against the following mouse antigens: Ly76 (TER-119), Gr-1 (L50-823), CD19 (KMC8), c-Kit (2B8), Sca-1 (D7), F4/80 (BM8), Gr-1 (RB6-8C5), CD45 (30F11), Flk1 (Avas12a1), CD31 (MEC13.3).

### Quantitative real-time PCR and analysis on mouse fetal liver cell subsets

Cells were sorted in RLT Buffer (Qiagen) and were frozen at −80 °C. RNA was extracted using the RNeasy Micro Kit (Qiagen), and cDNA was obtained with PrimeScript^™^ RT Reagent Kit (Takara). A 7300 Real-Time PCR System (Applied Biosystem) and Taqman technology (Applied Biosystem) were used for quantitative RT-PCR. Statistical analysis was completed using one-way ANOVA, multiple comparison test. All primers used are listed in Supplemental Table2.

