## supplemental files - tables and figures for "Myeloid and endothelial cells cooperate to promote hematopoietic stem cells expansion in the fetal niche"

Supplemental Table 1. Primer used for quantitative real time PCR of zebrafish expressed genes.

| | |
|---|---|
| ptger 4a-F | TGCCAATATTTCGGCTTCGTGCTG |
| ptger 4a-R | ATGCGTAAATGGCGAGTAGGGTGA |
| ptger1a-F | ACCTGGTGCAATAGTCATGAGGCT |
| ptger1a-R | AGAAAGAGCGGACACAGTCCGAAA |
| ptger2a-F | TGCGGATACATCACCATCCCTTGT |
| ptger2a-R | GTGGCGTAAACATTGGCATACGCT |
| ptger3-F | TTATTCAGTTGATGGGCATTATGT |
| ptger3-R | AATTACAGTCCTTCTGCAATTCCT |
| cox1-F | ACAGTTCCAGTACCAGAACCGCAT |
| cox1-R | TCCACCAGCTTCTCCAAGCCATAA |
| cox2a-F | CACTGTTGCCGGACAACTTTCAGA |
| cox2a-R | TCCAGCAGTCTGTTTGGTGAAGGA |
| cox2b-F | CTTTACCATTGGCACCCCCT |
| cox2b-R | CCACCCTTAACACTGCTGGT |
| ptges3a-F | TATTGGAGGCCATTGACCC |
| ptges3a-R | GGACAATTCTTCATCCGAGTC |
| ptges3b-F | CTCAGTGGAGCAGATAATGT |
| ptges3b-R | GCTCCGTCTAGATCAGGTAA |
| pla2g4aa-F | CTCTCCCTTCAGCGGCATCA |
| pla2g4aa-R | GGTCAAGCCACTGTCCACCA |
| pla2g4ab-F | CATCAATCCAGAGTGGAATGAG |
| pla2g4ab-R | CCTATTAGGAATGACACCAGC |
| slco2b1-F | TTGCCCTGCCTCACTTCATT |
| slco2b1-R | AGGCTGGAGTTGAGTCTGGT |
| abcc4-F | GTCCGCCTCACCGTCACTC |
| abcc4-R | CGGCTCTTTCTTCTCCTCCTG |

Supplemental Table 2. Primer used for quantitative real time PCR of mouse expressed genes.

| | |
|---|---|
| Mouseq-Pla2g4a-F | CTACGTGCCACCAAAGTAAC |
| Mouseq-Pla2g4a-R | CCTAGGGTTTCATCCATGAC |
| Mouseq-Pla2g4b-F | CCCTGTCTGGAATCAGAACT |
| Mouseq-Pla2g4b-R | CTGATGAGCTGTTCCTCACA |
| Mouseq-Pla2g4c-F | GCTCCAGTCATTGCTGTCTT |
| Mouseq-Pla2g4c-R | CTCCAGGCTCTCATGAAAGT |
| Mouseq-Cox1-F | CCAGTGTGATTGTACTCGCA |
| Mouseq-Cox1-R | GTAGTCATGCGCTGATGGTG |
| Mouseq-Cox2-F | CGGACTGGATTCTATGGTGA |
| Mouseq-Cox2-R | GTGCACATTGTAAGTAGGTGG |
| Mouseq-Slco2b1-F | CTCAGGACTCACATCAGGAT |
| Mouseq-Slco2b1-R | GCTGCCGAAATAGCTCACAA |
| Mouseq-Ptges3-F | CAGCCTGCTTCTGCAAAG |
| Mouseq-Ptges3-R | GAAGTCCACACTGAGCCA |
| Mouseq-Abcc4-F | CAGAAGATCGCTCAAAGCAC |
| Mouseq-Abcc4-R | CTGCACGTGGTAGAAGTACA |
| Mouseq-PTGER1-F | TTTATTAGCCTTGGGCCTCG |
| Mouseq-PTGER1-R | TTGCACACTAATGCCGCAAG |
| Mouseq- PTGER2-F | GAATTGGTGCTCACTGACCT |
| Mouseq-PTGER2-R | CATCGTGGCCAGACTAAAGA |
| Mouseq-PTGER3-F | TGTCGGTTGAGCAATGCAAG |
| Mouseq-PTGER3-R | GCAGAACTTCCGAAGAAGGA |
| Mouseq-PTGER4-F | CATCTTACTCATCGCCACC |
| Mouseq PTGER4-R | GCACAGTCTTCCGAAGAAG |

#### Supplemental Figures

**Suppl. Fig. 1 The prostaglandin transporters *slco2b1* and *abcc4* are expressed in the CHT niche.**

(a) WISH for *slcob1* expression during zebrafish embryo development at different stages. (b) WISH for *abcc4* expression during zebrafish embryo development at 48hpf. Scale bar 500μm (a-b).

**Suppl. Fig. 2 A new zebrafish model for myeloid ablation**.

(a) the *cd45:CFP-NTR* transgenic construct. (b) Cytometry performed on double transgenic embryos *cd45:CFP-NTR/runx1:mcherry* at 72hpf.

**Suppl. Fig. 3 Myeloid ablation decreases PGE2 level in the CHT.**

(a) Fluorescence imaging of *cd45:CFP-NTR* embryo in DMSO and after 24 hours of MTZ treatment. (b) Experimental outline to measure PGE2 level by ELISA-kit in 72hpf *cd45:CFP-NTR* embryos after 24 hours of DMSO or MTZ treatment. Quantification of PGE2. The statistical analysis was completed using an unpaired two tailed t test ***P < .001. Scale bar 500μm (a)

**Suppl.Fig.4 qPCR screening of PGE2 synthesis pathway after myeloid ablation.**

(a) Experimental outline of qPCR analysis after myeloid ablation in *cd45:CFP-NTR* embryos by MTZ treatment, to measure the expression of PGE2 synthesis gene pathway. (b-c) The expression of phospholipases *pla2g4aa* and *pla2g4ab* was highly reduced after myeloid ablation. (d-e-f) So is the expression of cyclooxygenases *cox1, cox2a and cox2b*. (g-h-i-l) No difference of expression for the prostaglandin transporters *slco2b1* and *abcc4*, nor for the prostaglandin synthase *ptges3a*. *ptges3b* was midly reduced after myeloid ablation. (m-n-o-p-q) Significative reduction for the different PGE2 receptors after myeloid ablation.

**Suppl.Fig.5 The loss of HSCs after myeloid ablation cannot be rescued by AA or PGG2 treatments.**

(a) Fluorescence imaging in the CHT of double transgenic *cd45:CFP-NTR/runx1:mcherry* embryos in DMSO and after treatment with MTZ and AA or PGG2. (b) Quantification of *runx1:mcherry* positive-cells in double transgenic *cd45:CFP-NTR/runx1:mcherry* embryos in DMSO and after treatment with MTZ and AA or PGG2. (c) Quantification of *cd45:CFP* positive-cells in double transgenic *cd45:CFP-NTR/runx1:mcherry* embryos in DMSO and after treatment with MTZ and AA or PGG2. Statistical analysis was completed using one-way ANOVA, multiple comparison test. **P<.01; ****P<.0001. Scale bar 200µm (a).

**Suppl. Fig. 6 *Slco2b1^-/-^* mutants present a decrease of HSPCs in the CHT.**

(a) Sequencing of PCR product to genotype the *slco2b1^sa37367^* mutant line. This mutant presents a point mutation T > C in the splice donor site at the end of exon 4. (b) WISH for *cmyb* expression at 4dpf in sibling and *slco2b1^-/-^*. (c) WISH for *runx1* expression at 28hpf in sibling and *slco2b1^-/-^*.(d) WISH for *cmyb* expression at 36hpf in sibling and *slco2b1^-/-^*. Scale bar 500µm (b); 200µm (c-d).

**Suppl. Fig. 7 *Slco2b1*-deficiency does not affect primitive erythropoiesis, vasculogenesis and myelopoiesis.**

(a) WISH for *kdrl* at 24hpf in sibling and *slco2b1^-/-^*embryos. (b) WISH for *gata1* at 22hpf in sibling and *slco2b1 ^-/-^* embryos. (c) WISH for *pu.1* at 22hpf in sibling and *slco2b1^-/-^* embryos. Scale bar 500µm (a-b-c).

**Suppl. Fig. 8 Morpholino-mediated knockdown of *slco2b1* expression fully phenocopies *slco2b1^-/-^* mutants.**

(a) Schematic of MO-retention targeting intron/exon junctions in *slco2b1*. Validation of the *slco2b1*-MO that induces intron-3 retention. The RT-PCR was performed on mRNA/cDNA obtained at 48hpf from pools of 10 embryos. The PCR products were sequenced using the forward primer. (b) Wish for *runx1* expression at 28hpf in control and *slco2b1*-morphant. (c) WISH for *cmyb* expression at 36hpf control and *slco2b1*-morphant. (d) WISH for *cmyb* expression at 4dpf control and *slco2b1*-morphant. Scale bar 200µm (b-c); 500µm (d).

**Suppl. Fig. 9 The loss of HSPCs in *slco2b1*-deficient embryos can only be rescued by PGE2 treatment.**

(a) WISH for *cmyb* expression at 60hpf in embryos injected with control- or *slco2b1*-morpholinos, and treated with DMSO and/or AA, PGG2, PGH2, PGE2. (b) Quantification of the number of *cmyb*-expressing cells. Statistical analysis was completed using one-way ANOVA, multiple comparison test. *P<.05; **P<.01; ****P<.0001. (c) WISH for *cmyb* expression at 60hpf in siblings and *slco2b1^-/-^* embryos, either non-treated or after PGE2 supplementation. (d)

Quantification of the number of *cmyb*-expressing cells. The statistical analysis was completed using an unpaired two tailed t test *P < .01; ***P < .001; ****P < .0001. Scale bar 200µm (a-c).

**Suppl. Fig. 10 *Slco2b1*-overexpression in ECs or macrophages through the Gal4:UAS system**.

(a) WISH for *slco2b1* at 60hpf in *kdrl:Gal4*+ embryos non-injected or after co-injection with the Tol2-UAS:*slco2b1* vector and *tol2* mRNA. (b) WISH for *slco2b1* at 60hpf in *mpeg1:Gal4*+ embryos non-injected and after or after co-injection with the Tol2-UAS:*slco2b1* vector and *tol2* mRNA.

**Suppl. Fig. 11 *Slco2b1*-overexpression in ECs can rescue the loss of HSPCs in *slco2b1*-morphants.**

(a) WISH for *cmyb* at 60hpf in *kdrl:Gal4*+ embryos injected with control and *slco2b1*-morpholinos and/or co-injected with the Tol2-UAS:*slco2b1* vector and *tol2* mRNA. (b) magnification of CHT regions. (c) Quantification and statistical analysis were completed using one-way ANOVA, multiple comparison test. ****P<.0001.

**Suppl. Fig. 12 *Slco2b1*-overexpression in macrophages cannot rescue the loss of HSPCs in *slco2b1*-morphants.**

(a) WISH for *cmyb* at 60hpf in *mpeg1:Gal4*+ embryos injected with control and *slco2b1*-morpholinos and/or co-injected with the Tol2-UAS:*slco2b1* vector and *tol2* mRNA. (b) magnification of CHT regions. (c) Quantification and statistical analysis were completed using one-way ANOVA, multiple comparison test. ****P<.0001.

**Suppl. Fig. 13 The PGE2 synthesis pathway is conserved in the mouse fetal liver.**

qPCR analysis on FACS-sorted cells from mouse fetal livers at Embryonic day (E) 13.5 by using various combinations of antibodies. HSCs were isolated based on their Lin-Sca1+cKit+ (LSK) phenotype; F4/80 to sort macrophages, *Flk1* to mark ECs and Gr-1, known as Ly-6G/Ly-6C, to sort neutrophils. (a) The phospholipases *pla2g4a* are highly expressed in neutrophils respect to (b-c) *pla2g4b* and *pla2g4c*. (d-e) The expression of cyclooxygenases *cox1* and *cox2* is enriched in macrophages. (f-h) The prostaglandin synthases *ptges3* and the prostaglandin transporters *slco2b1*, *abcc4* are specifically expressed in ECs. (i-n) The prostaglandin receptors *ptger1, ptger4* are mostly expressed in HSPCs respect to *ptger2, ptger3.* Statistical analysis was completed using one-way ANOVA, multiple comparison test. *P<.05; **P<.01; ***P<.001; ****P<.0001.

**Suppl.Movie 1. Time-lapse confocal imaging of the CHT of a *cmyb:GFP* embryo injected with control morpholino (ctrl-mo). (54-60hpf)**

**Suppl.Movie 2. Time-lapse confocal imaging of the CHT of a *cmyb:GFP* embryo injected with slco2b1-morpholino (slco2b1-mo). (54-60hpf)**

### Supplementary figure 1

**a**

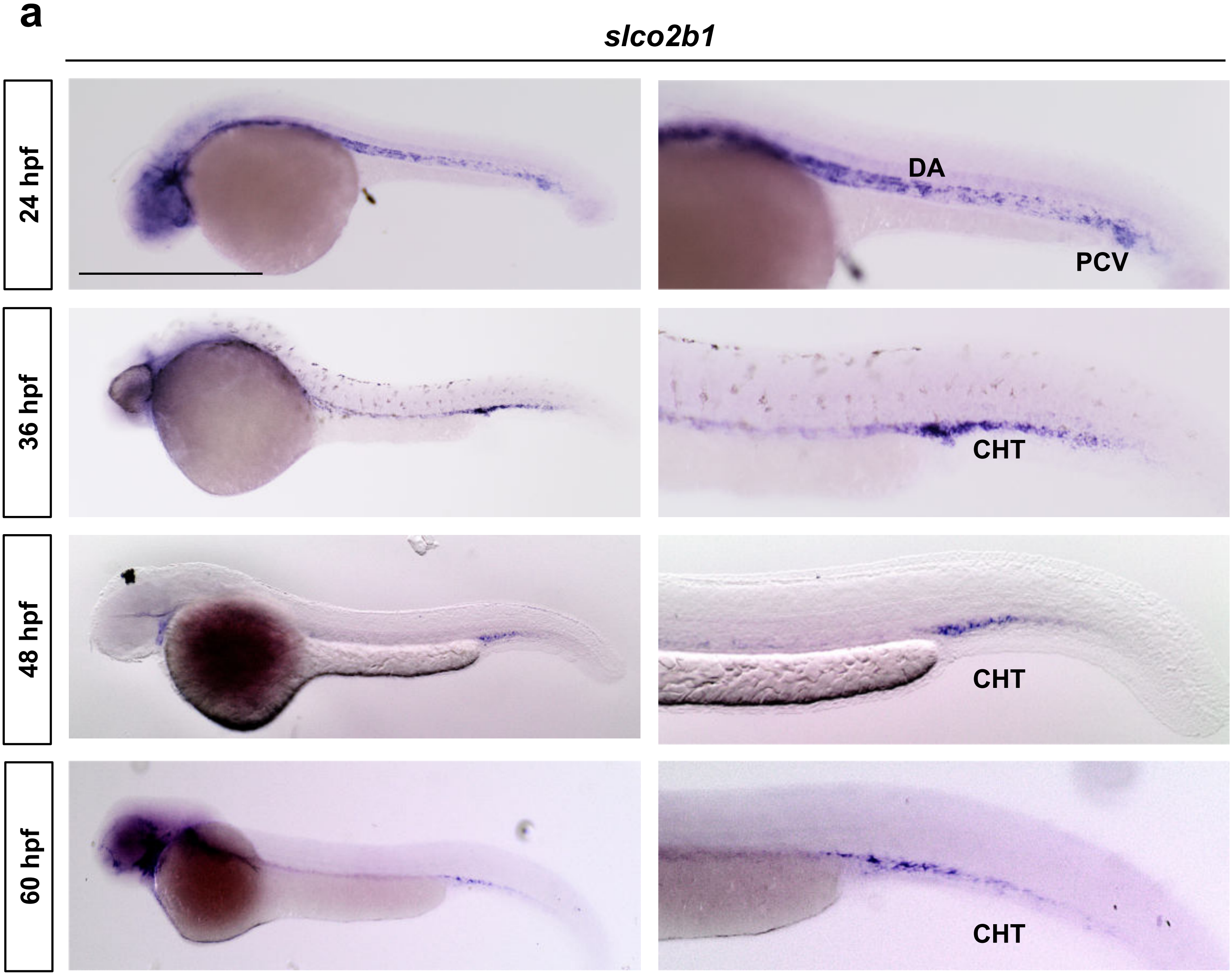


**b**

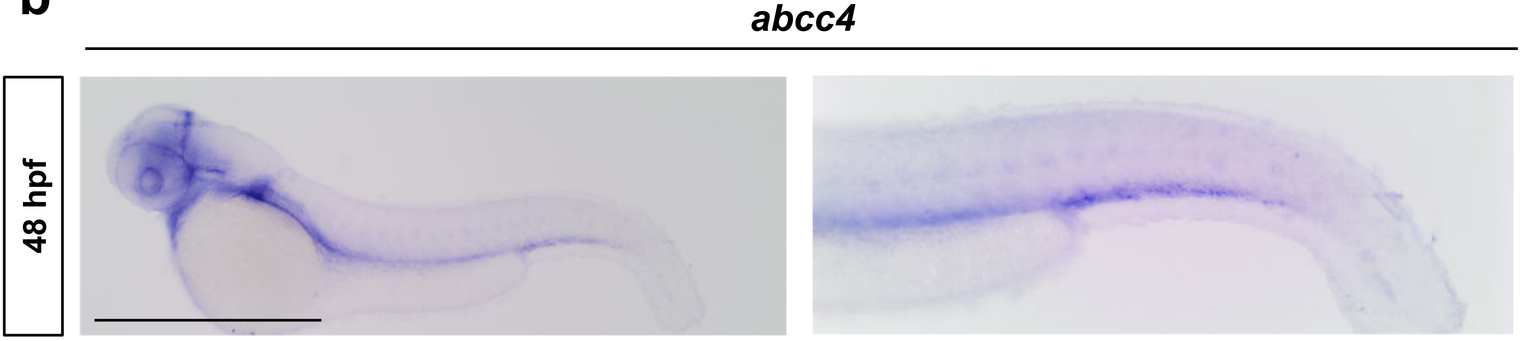

#### Supplementary figure 2

**a**

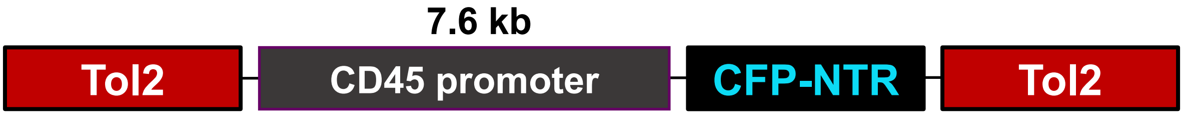


**b**

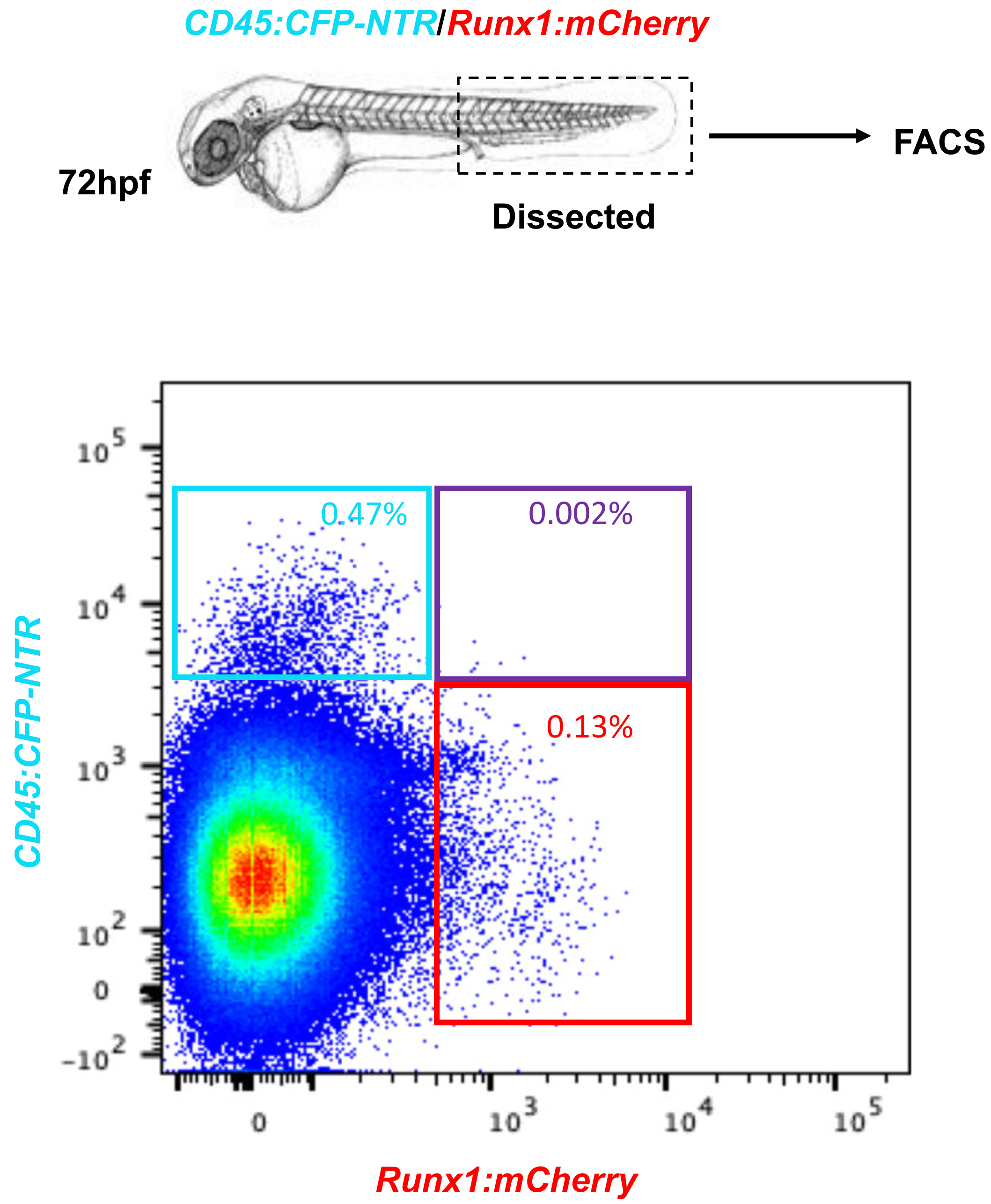

### Supplementary figure 3

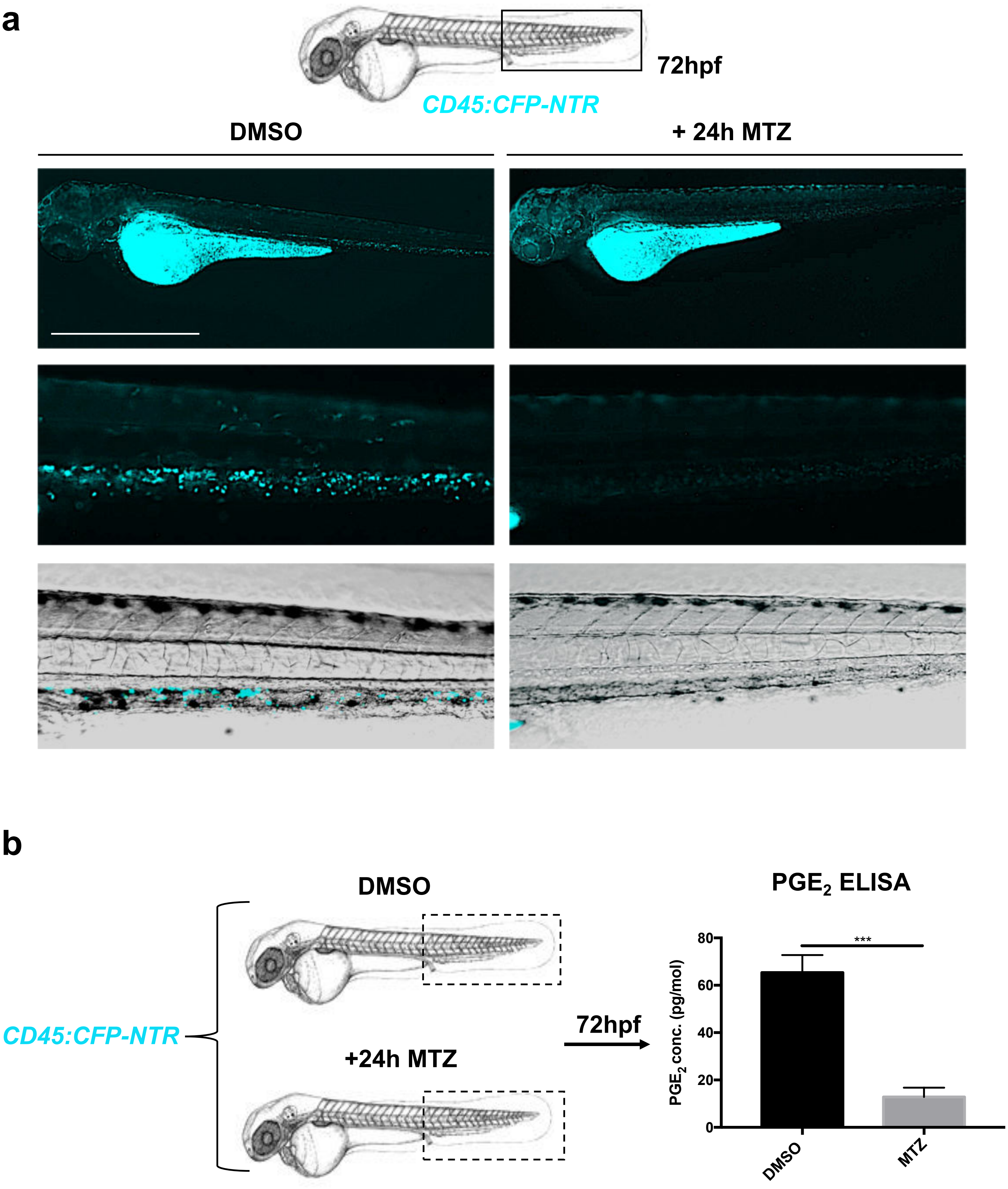

### Supplementary figure 4

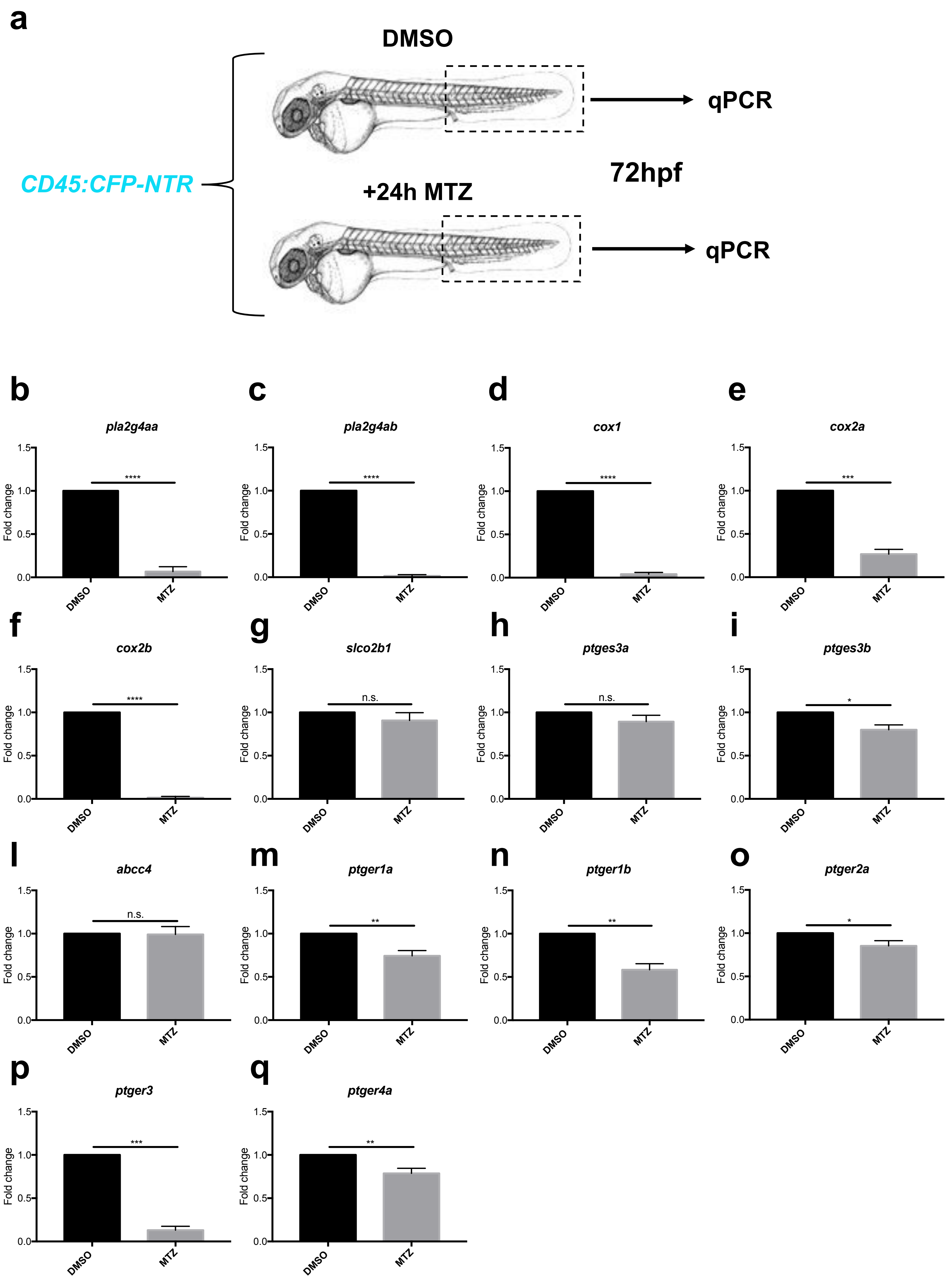

### Supplementary figure 5

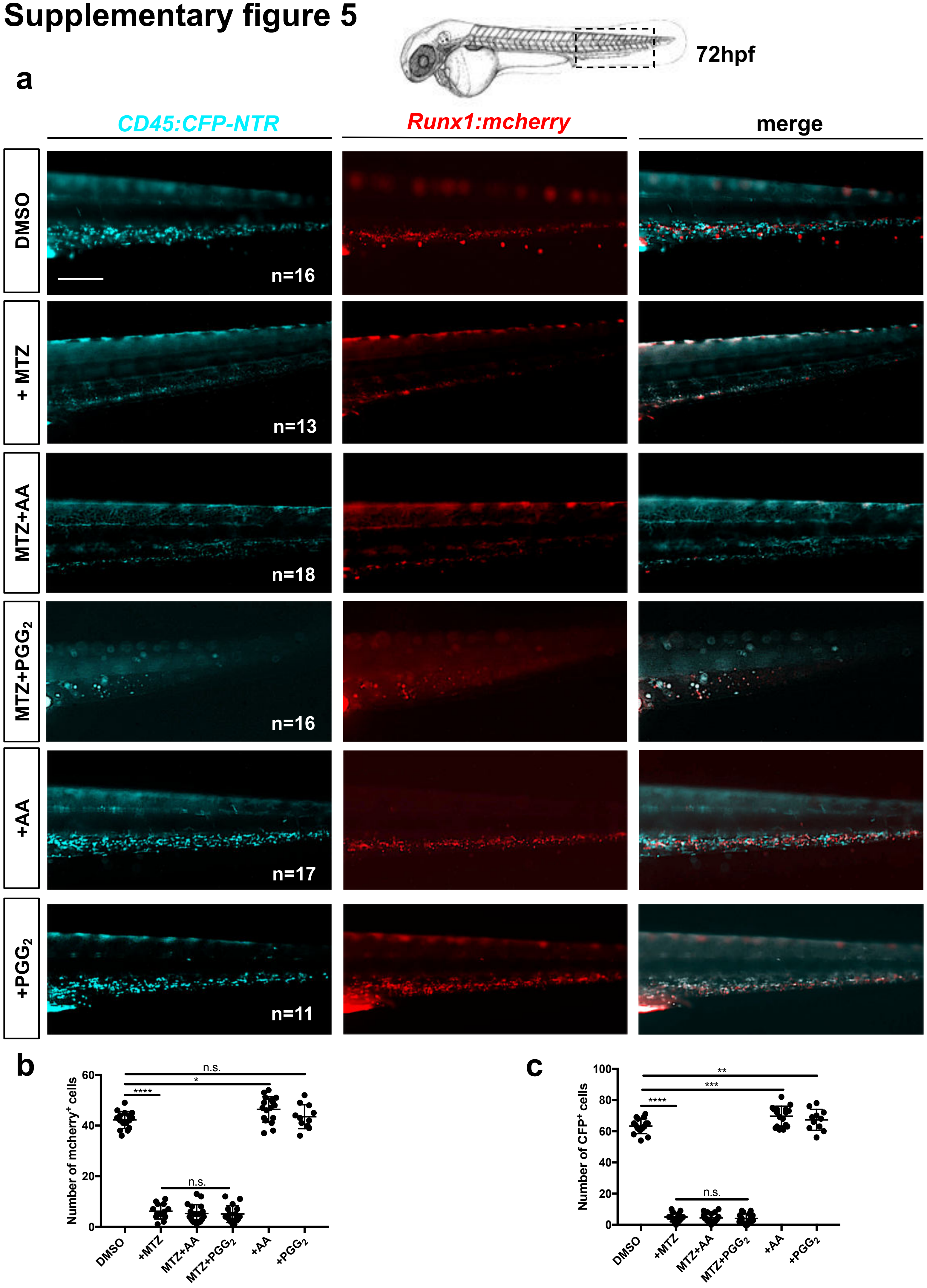

### Supplementary figure 6

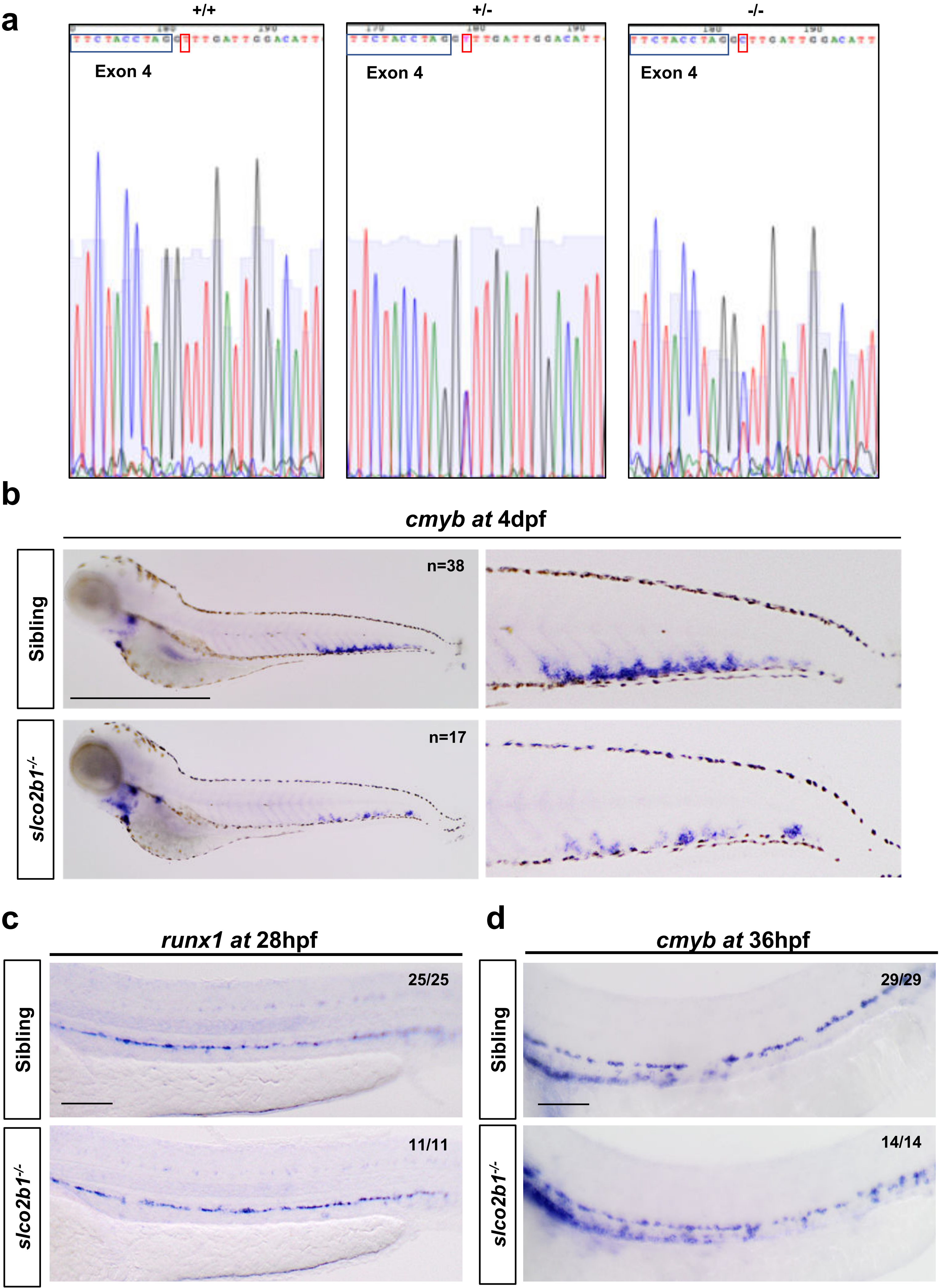

### Supplementary figure 7

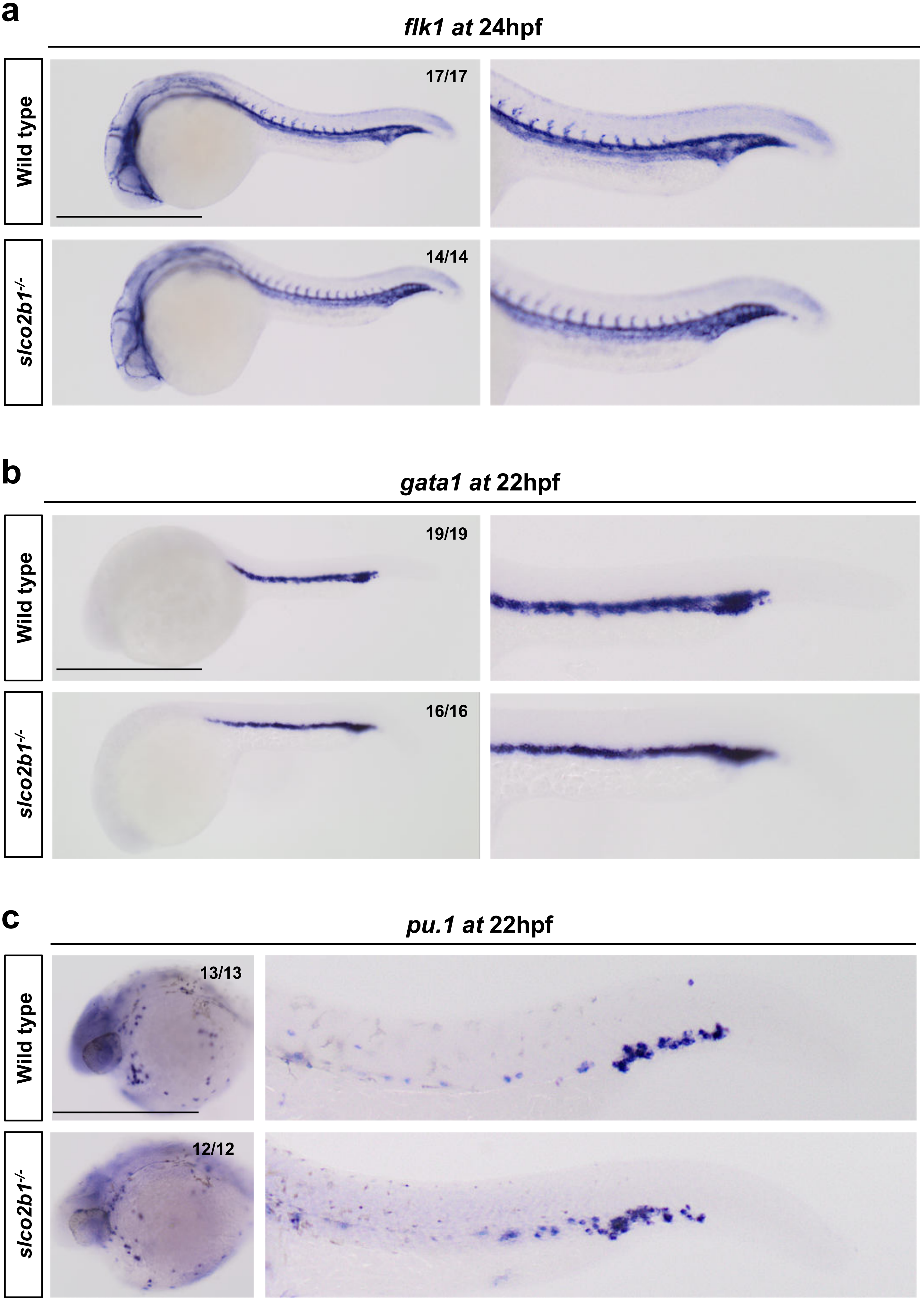

### Supplementary figure 8

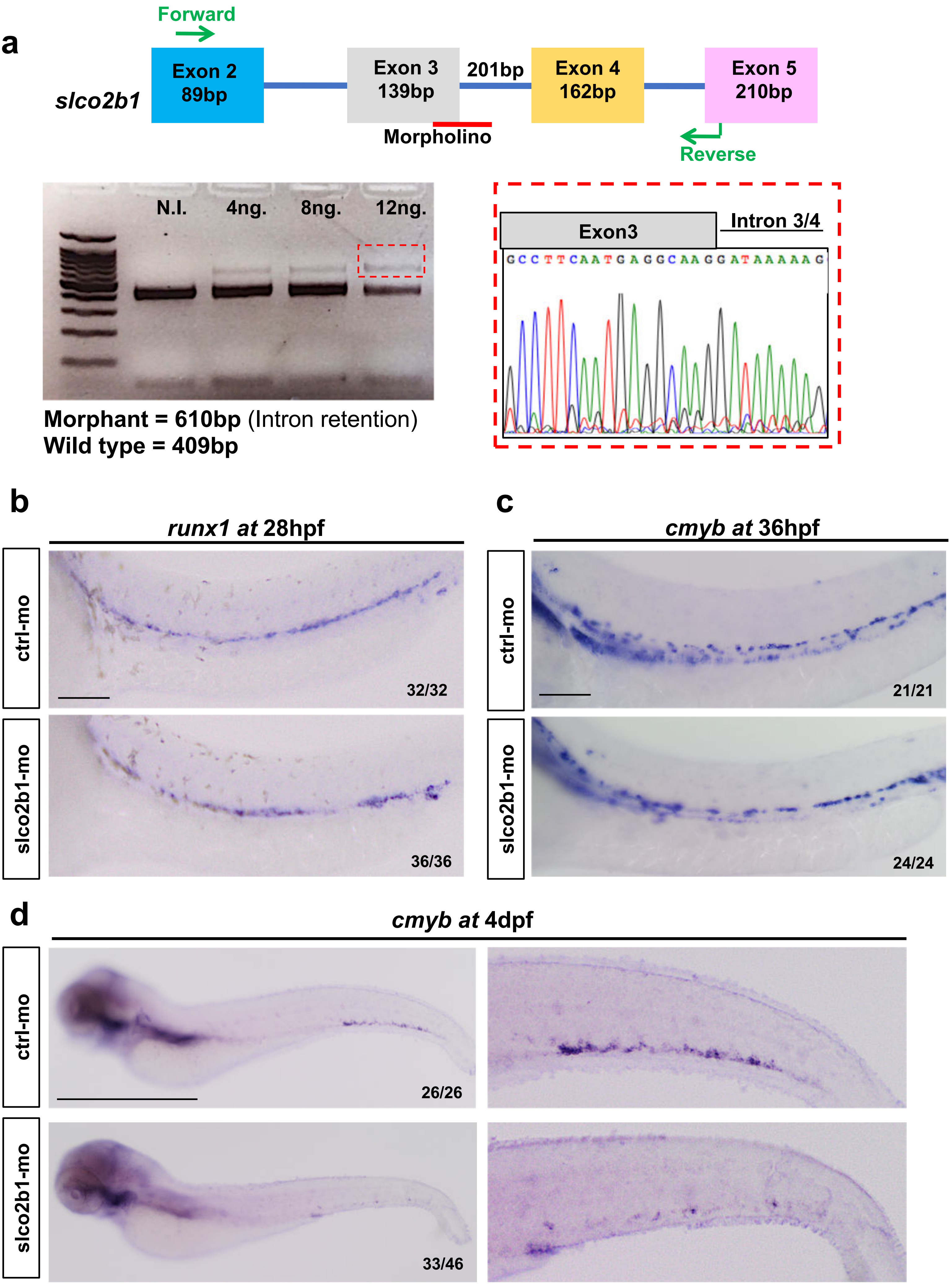

### Supplementary figure 9

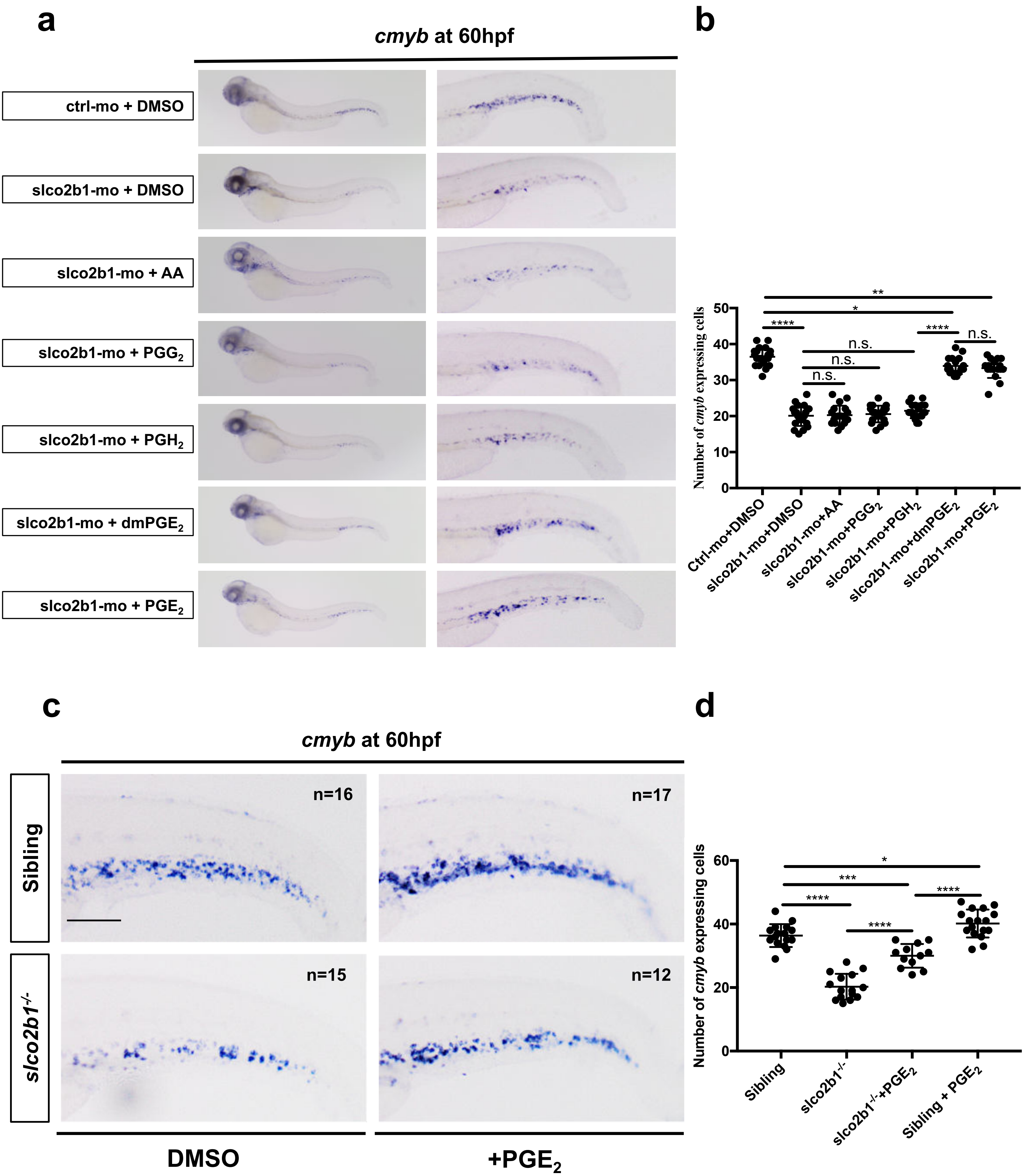

### Supplementary figure 10

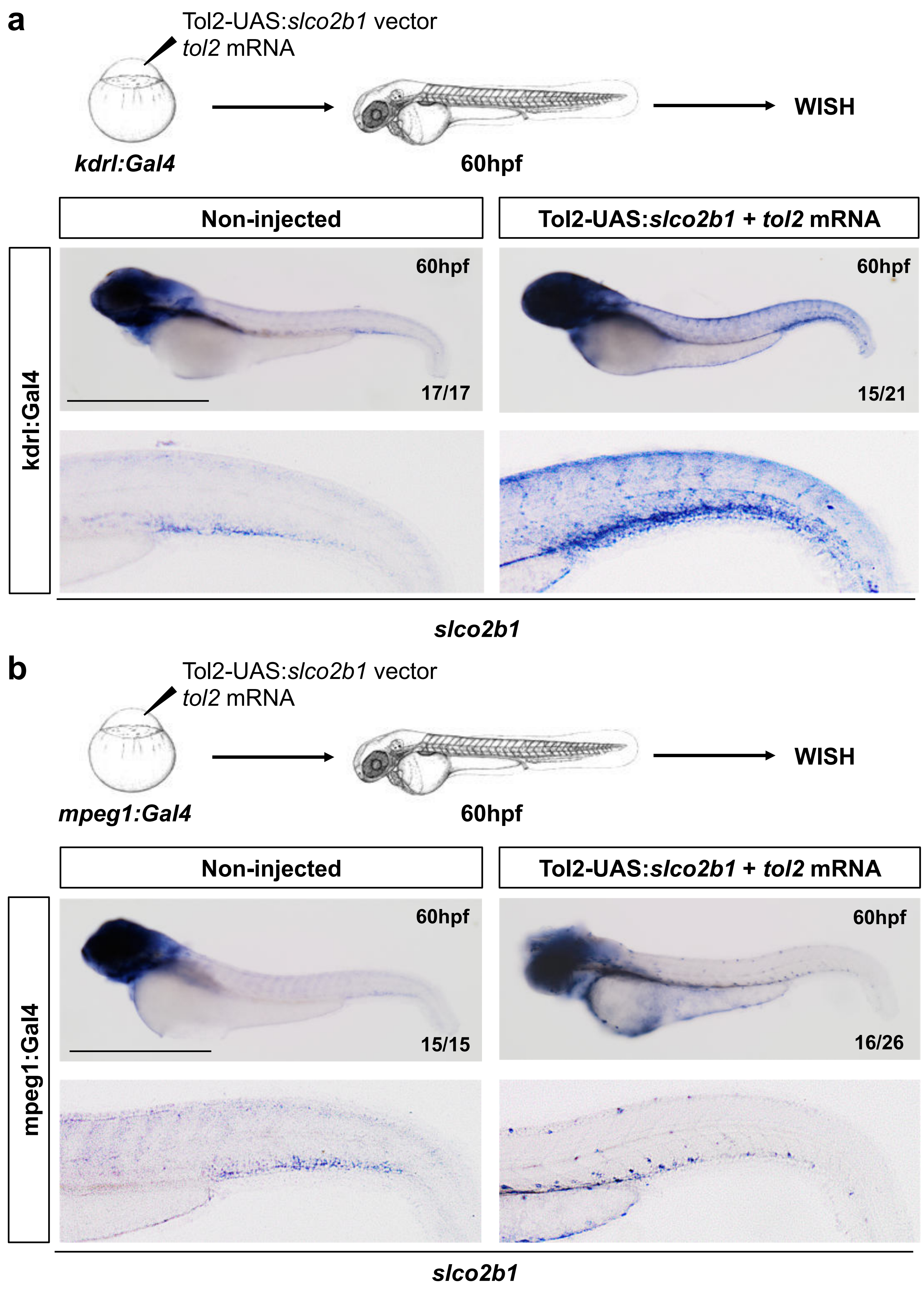

### Supplementary figure 11

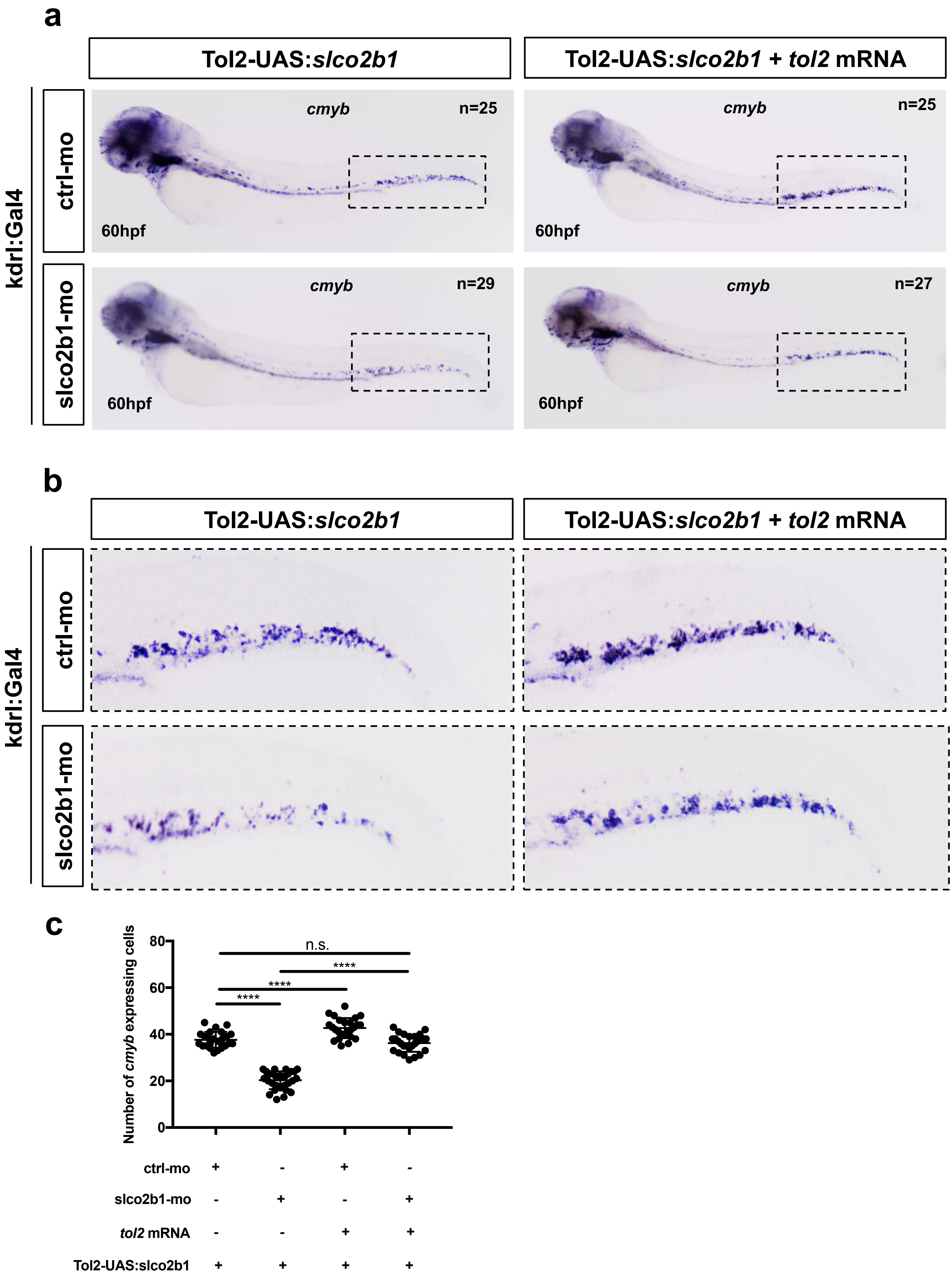

### Supplementary figure 12

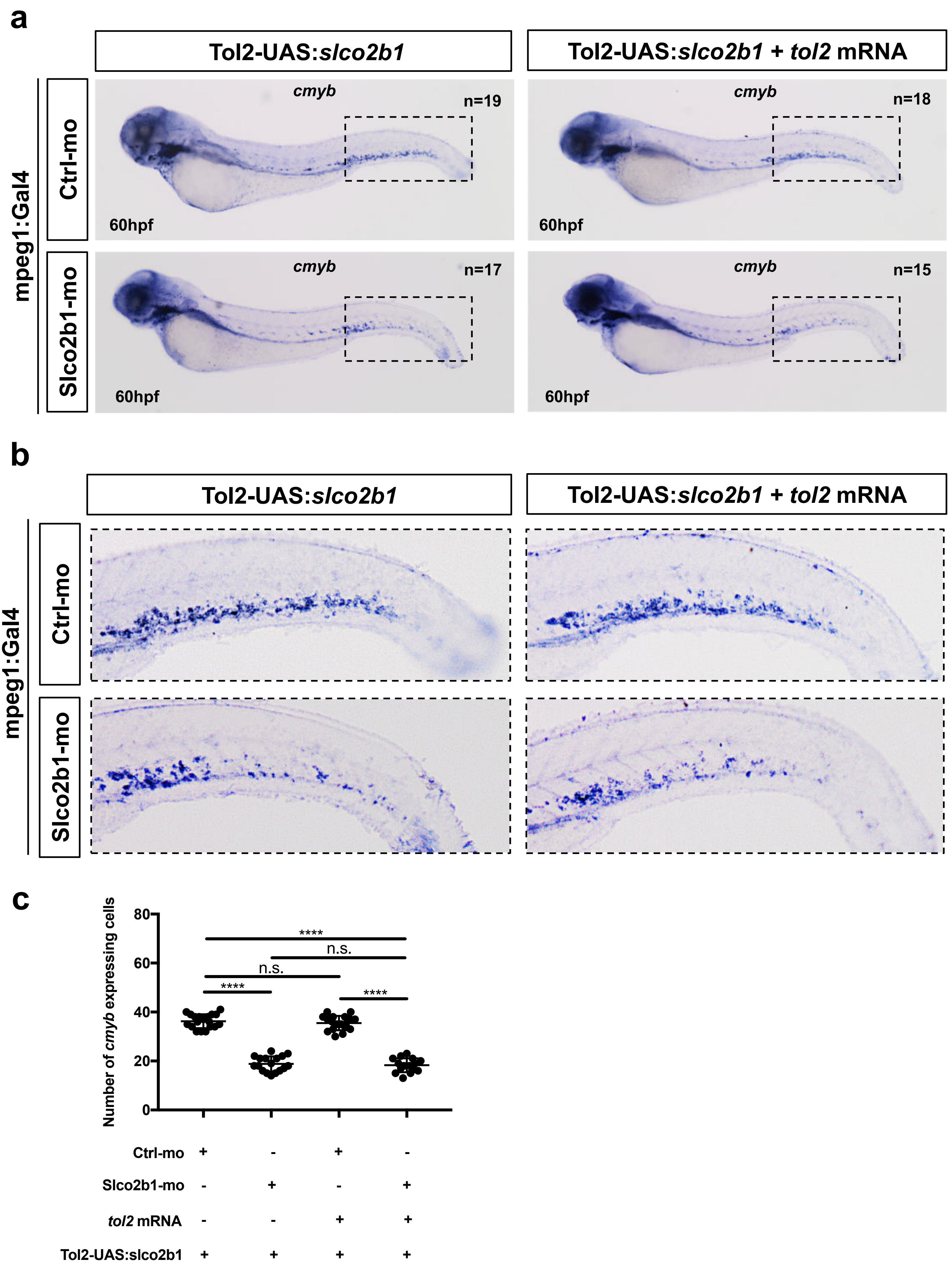

### Supplementary figure 13

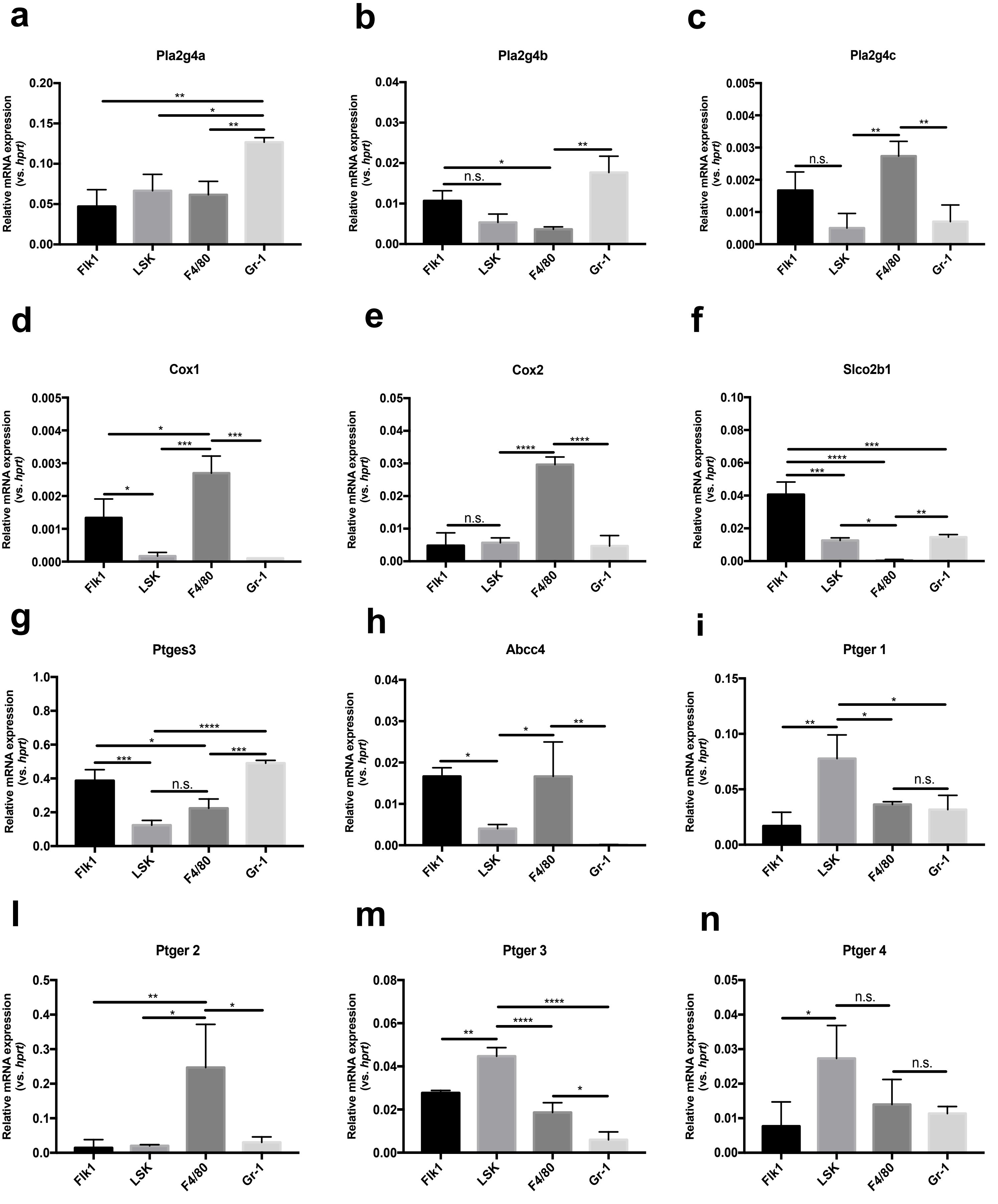
